# Human brain-derived Aβ oligomers bind to synapses and disrupt synaptic activity in a manner that requires APP

**DOI:** 10.1101/135673

**Authors:** Zemin Wang, Rosemary J. Jackson, Wei Hong, Taylor M. Walter, Arturo Moreno, Wen Liu, Shaomin Li, Matthew P. Frosch, Inna Slutsky, Tracy Young-Pearse, Tara L. Spires-Jones, Dominic M. Walsh

## Abstract

Compelling genetic evidence links the amyloid precursor protein (APP) to Alzheimer’s disease (AD), and several theories have been advanced to explain the involvement of APP in AD. A leading hypothesis proposes that a small amphipathic fragment of APP, the amyloid β-protein (Aβ), self-associates to form soluble aggregates which impair synaptic and network activity. Here, we report on the plasticity-disrupting effects of Aβ isolated from AD brain and the requirement of APP for these effects. We show that Aβ-containing AD brain extracts block hippocampal long-term potentiation (LTP), augment glutamate release probability and disrupt the excitation/inhibition balance. Notably, these effects are associated with Aβ localizing to synapses, and genetic ablation of APP prevents both Aβ binding and Aβ-mediated synaptic dysfunctions. These findings indicate a role for APP in AD pathogenesis beyond the generation of Aβ and suggest modulation of APP expression as a therapy for AD.

**Acknowledgments:** We thank Dr. Tiernan T. O’Malley for useful discussions and technical advice. This work was supported by grants to DMW from the National Institutes of Health (AG046275), Bright Focus, and the United States-Israel Binational Science Foundation (2013244, DMW and IS); grants to TSJ from Alzheimer’s Research UK and the Scottish Government (ARUK-SPG2013-1), Wellcome Trust-University of Edinburgh Institutional Strategic Support funds, and the H2020 European Research Council (ALZSYN); and to the Massachusetts Alzheimer’s Disease Research Center (AG05134).

## Introduction

Mutation, over-expression or altered-processing of the amyloid precursor protein (APP) underlie all known monogenic cases of familial Alzheimer’s disease (fAD) (Tanzi, 2012; Guerreiro and Hardy, 2014). Although the physiological roles of APP are not fully understood, myriad studies indicate that APP plays a role in synaptic plasticity, dendritic morphogenesis, and neuroprotection (Muller and Zheng, 2012). Membrane-tethered APP can act as a cell-adhesion molecule linking the pre-and post-synapse (Soba et al., 2005) and APP has been shown to regulate synaptic vesicle proteins, synaptic transmission and plasticity (Seabrook et al., 1999; Lassek et al., 2013; Fanutza et al., 2015; Lassek et al., 2016). In the dentate gyrus (DG) of rat, APP expression is known to change during memory consolidation (Conboy et al., 2005) and intraventricular administration of anti-APP antibodies or antisense oligonucleotides results in profound amnesia (Doyle et al., 1990; Huber et al., 1993; Mileusnic et al., 2000). Notably, APP is a component of the presynaptic GABA-B1a receptor (GABA_B1a_-R) complex (Bai et al., 2008; Schwenk et al., 2016) and neuron-type specific knock-out of APP indicates an important role for APP in GABAergic transmission and maintenance of the excitatory–inhibitory balance (Wang et al., 2014).

APP is a complex molecule that undergoes substantial post-translational modification and processing. More than 10 different proteolytic fragments of APP have been identified (Weidemann et al., 1989; Esch et al., 1990; Golde et al., 1992; Haass et al., 1992; Sisodia, 1992; Portelius et al., 2013; Welzel et al., 2014; Willem et al., 2015). Several of these are suggested to be pathogenic (Neve and McPhie, 2007; Yankner and Lu, 2009; Tamayev et al., 2012; Willem et al., 2015), whereas others are neuroprotective (Mockett et al., 2017). The fragment from which the precursor protein derives its name, the amyloid β-protein (Aβ), is found in the tell-tale amyloid plaques which litter the brains of individuals who die with AD. Aβ comprises a family of APP-derived peptides that share a common core of ∼30 amino acids (Walsh and Teplow, 2012) which are produced by the concerted action of two aspartyl proteases, β–secretase and γ-secretase (De Strooper, 2010). Aβ peptides are prone to self-associate and multiple studies indicate that certain forms of Aβ adversely affect synaptic form and function (Shankar and Walsh, 2009).

The synaptotoxic activity of Aβ and the involvement of APP in synapse formation and activity are particularly relevant to AD since *in vivo* and postmortem studies indicate that synapse dysfunction and loss are prominent early features of AD (Scheff et al., 2006; Scheff et al., 2007; Johnson et al., 2012). Transgenic (tg) mice over-expressing APP either alone, or in combination with PS1, produce high levels of Aβ, deposit amyloid plaques, and exhibit deficits in learning and memory (Ashe and Zahs, 2010). Certain APP tgs also manifest aberrant changes in synapses, neuronal microcircuits and complex networks (Palop and Mucke, 2016) and some authors have sought to link the network hyperactivity observed in APP tgs with epileptiform changes detected in a segment of individuals with early stage AD (Palop and Mucke, 2009; Busche and Konnerth, 2015). However, there is no human parallel for the high level of APP over-expression seen in APP tg mice. This supraphysiological production of APP induces artifacts such as increased mortality and behavioral hyperactivity (Ashe and Zahs, 2010; Nilsson et al., 2014), and it is difficult to differentiate between effects mediated by Aβ, APP, or non-Aβ APP derivatives (Seabrook et al., 1999). Indeed, there is now evidence that the epileptiform changes seen in APP tgs, that had formerly been attributed to Aβ, are in fact mediated by non-Aβ APP derivatives (Born et al., 2014). Surprisingly, mice which produce and deposit human Aβ, but do not over-express human APP (i.e. APP knock-in mice or BRI2-Aβ mice) show no changes in electroencephalogram (EEG) activity or deficits in synaptic plasticity (Kim et al., 2013; Born et al., 2014).

Acute studies in wild type rodents show that non-fibrillar, water-soluble Aβ from a variety of sources are potent synaptotoxins (Lambert et al., 1998; Walsh et al., 2002; Cleary et al., 2005; Klyubin et al., 2008; Minkeviciene et al., 2009; Kurudenkandy et al., 2014). Furthermore, *in vitro* and *in vivo* studies demonstrate that the most disease-relevant form of non-fibrillar Aβ, Aβ extracted from the water-soluble phase of AD brain, inhibits long-term potentiation (LTP), facilitates long-term depression (LTD), reduces synaptic remodeling, and impairs memory consolidation (Shankar et al., 2008; Barry et al., 2011; Freir et al., 2011; Borlikova et al., 2013; Hu et al., 2014; Yang et al., 2017). Here, we show that the block of LTP mediated by Aβ-containing AD brain extracts is accompanied by opposing changes in excitatory and inhibitory pre-synaptic release probabilities and consequent disruption of the excitation/inhibition (E/I) balance. The net increase in the E/I ratio and inhibition of LTP require expression of APP and are associated with Aβ localizing to synapses. These findings suggest a link between Aβ toxicity and perturbation of the normal regulatory role of APP, and are consistent with prior studies which have imputed a role for APP in Aβ toxicity (White et al., 1998; Lorenzo et al., 2000; Shaked et al., 2006; Sola Vigo et al., 2009; Fogel et al., 2014; Kirouac et al., 2017). In light of these results we suggest that down-regulation of APP expression or modulation of its interaction with synaptotoxic Aβ species should be investigated as an approach to treat AD.

## Results

We previously reported that aqueous extracts of certain end-stage AD brains block hippocampal LTP *in vivo* and *in vitro* (Shankar et al., 2008; Li et al., 2009; Barry et al., 2011; Freir et al., 2011; Hu et al., 2014). Here we further investigated the mechanism of this effect and the requirement of endogenous APP.

### The water-soluble extract from AD brain contains both Aβ monomers and oligomers and blocks LTP in a manner dependent on Aβ

Brain extracts were prepared as described and a portion was immunodepleted (ID) of Aβ or mock-ID with pre-immune rabbit serum. Here, the mock-ID extract is referred to as the AD sample, and the material depleted of Aβ as ID-AD. ID-AD and AD samples were analyzed using IP/WB, and MSD immunoassays that preferentially recognize either Aβ oligomers (oAssay) or Aβ42 monomers (Mc Donald et al., 2015). IP/WB analysis allows the capture of Aβ structures under native conditions and their detection following denaturing SDS-PAGE. The Aβ present in the AD sample migrated on SDS-PAGE with molecular weights of ∼4 and ∼7-8 KDa (Figure 1A) — a pattern we have seen in aqueous extracts of more than 100 AD brains analyzed in our laboratory (Mc Donald et al., 2015). Since SDS-PAGE is highly denaturing, the ∼4 and ∼7 kDa species do not necessarily reflect native Aβ species. Rather, these simply indicate that at least two different Aβ species are present. The same samples were treated plus or minus 5 M GuHCl and then analyzed using MSD assays. In prior studies we found that GuHCl effectively disaggregates high molecular weight Aβ species such that the signal detected by our oAssay is greatly decreased, whereas the signal detected by the monomer-preferring Aβx-42 immunoassay is proportionately increased (Mc Donald et al., 2015). A similar outcome was evident when the extract of AD1 was treated with GuHCl (Figure 1B). Specifically, GuHCl treatment caused a ∼70% decrease in the oligomer signal and a more than 8-fold increase in the monomer signal. Together these immunoassay and IP/WB results indicate that the majority of Aβ in the AD1 extract exist as labile aggregates made up of ∼4 kDa Aβ and ∼7 kDa Aβ. Importantly, AW7 ID effectively removed the large majority of the various Aβ species detected (Figure 1A and B). For instance, AW7 ID reduced the oligomer signal from 5.1 ± 0.03 ng/ml to 0.32 ± 0.12 ng/ml (Figure 1B, left panel) and monomer from 3.42 ± 0.03 ng/ml to 0.12 ± 0.04 ng/ml (Figure 1B, right panel).

**Figure 1:**
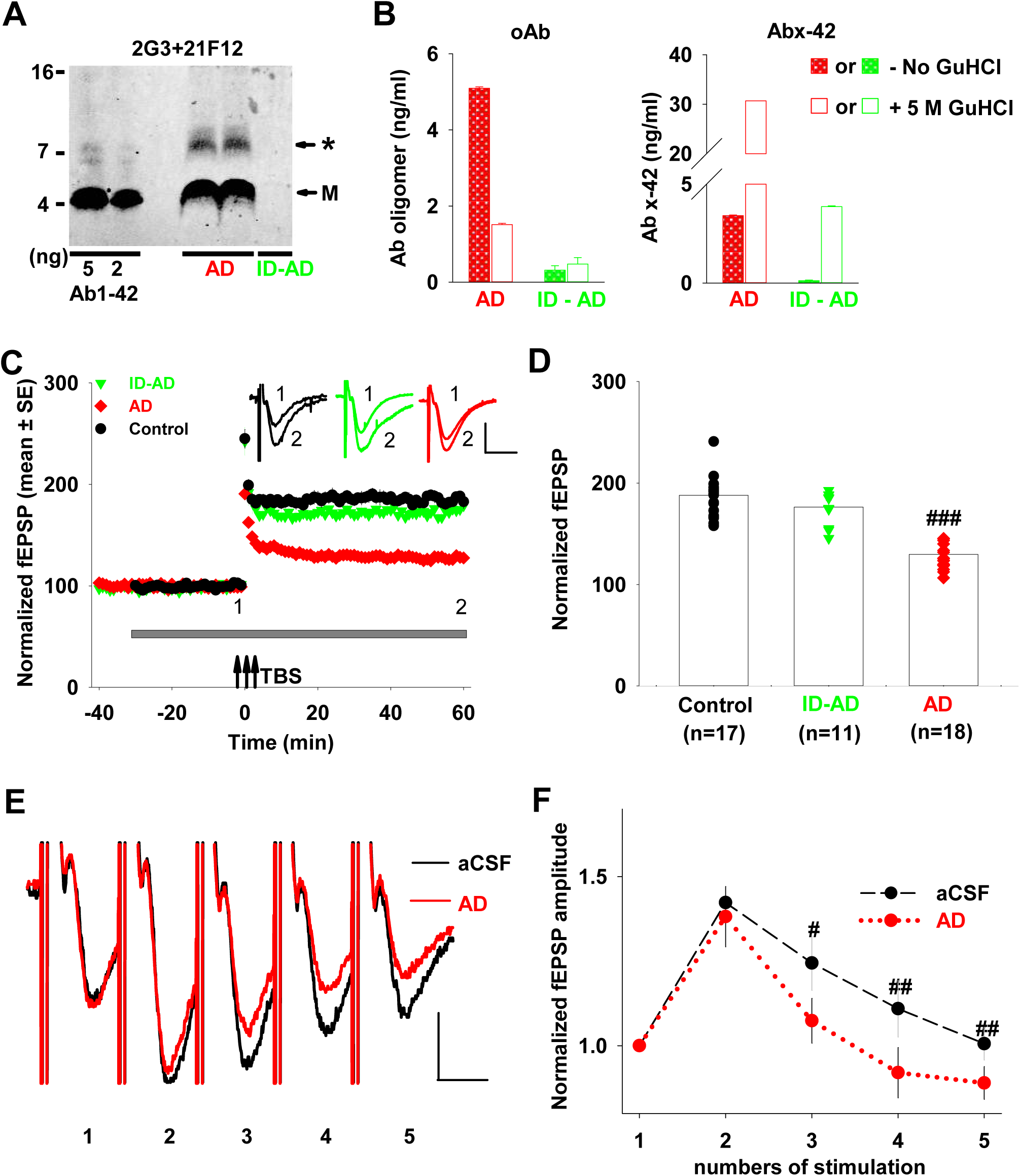
The water-soluble extract of AD brain contains both Aβ monomers and oligomers and perturbs short-term and long-term synaptic plasticity. (**A)** Aqueous extract of AD1 was treated with either pre-immune rabbit serum or with AW7 antiserum. Portions of the mock immunodepleted sample (AD, red) and the AW7 immunodepleted sample (ID-AD, green) were then analyzed by IP/WB, using AW7 for IP and a combination of 2G3 and 21F12 for WB. **M** denotes Aβ monomer and * indicates a broad smear ~7–8 kDa. No specific bands were detected above 16 kDa marker and the blot was cropped accordingly. (**B)** AD (red) and ID-AD (green) samples were incubated +/- 5 M GuHCl and analyzed using immunoassays that preferentially recognize Aβ oligomers (1C22-3D6b, left panel) or Aβ42 monomer (266-21F12b, right panel). Values shown are the mean ± SEM of triplicate measurements. **(C)** Time course plots show that the AD sample but not the ID-AD sample blocked hippocampal LTP. The gray horizontal bar indicates the time period when sample was present in the bath. 1, 2, indicate example traces from time points just prior to the theta burst stimulation (↑↑↑ TBS) (1) and 60 min after TBS (2), respectively. The aCSF control is shown using black circles; AD treatment is shown using red diamonds and ID-AD with green downward triangles. Each slice used for each treatment was from a different animal. Scale bar 0.2 mV, 10 ms. (**D)** Histogram plots of the average potentiation for the last 10 min of the traces shown in **C**. Treatment of slices with AD sample significantly inhibited LTP compared to the aCSF vehicle control (F=4.26, p=6.98E-9) and ID-AD treatment (F=4.14, p=3.56E-12); in contrast ID-AD had no effect on LTP relative to the vehicle control (F=4.23, p=0.16); One Way ANOVA test. Symbols are the same as in panel **C**. (**E**) Representative traces of averaged field recordings were collected after 5 stimulation bursts (inter-stimulation interval 20 ms, inter-burst interval 30 s) before (black, aCSF) and 30 min after perfusion with the AD sample (red). The trace shown for the AD samples are scaled so that the first response matches that of the aCSF control. Scale bars: 0.5 mV, 10 ms. (**F**) fEPSPs amplitude after 2 to 5 stimulations were normalized to the value obtained after the first stimulation. Compared to vehicle control, AD treatment induced a small but significant decrease in short-term synaptic facilitation, (p=0.02) after the 3rd, (p=0.004) the 4^th^ stimulation and 5^th^ stimulation (p=0.004); n = 6, student t-test. Values shown are the mean ± SEM. # p< 32 0.05; ## p< 0.01; ### p<0.001.

For slices that received vehicle aCSF-B (Control), TBS induced strong potentiation which lasted the whole recording period (Figure 1C, black dots, 181.1 ± 10.7 %, n = 17), and ID-AD allowed a similar response (green downward triangles, 173.6 ± 8.7 %, n = 11, p= 0.12, One Way ANOVA test) (Figure 1C and D). Consistent with prior reports (Shankar et al., 2008; Freir et al., 2011), application of the AD1 extract significantly decreased LTP compared to both the Control and ID-AD treatment (red diamonds, 136 ± 4.2 %, n = 18, F=4.26,p=6.98E-9 AD vs. Control; F=4.14, p=3.56E-12 AD vs. ID-AD, One Way ANOVA test). The fact that the ID-AD and AD samples are identical except that the latter contains more Aβ than the former, is evidence that some form of Aβ is responsible for the block of LTP induced by the AD1 extract.

### Aβ-containing AD brain extract affects presynaptic release probabilities

Accumulating evidence indicates that soluble Aβ species may interact with excitatory and inhibitory presynaptic terminals, modulate neurotransmitter release and cause synaptic dysfunction in the very early stages of AD (Nimmrich et al., 2008; Abramov et al., 2009; Kabogo et al., 2010; Parodi et al., 2010; Russell et al., 2012; Sokolow et al., 2012; Huang et al., 2013; Ripoli et al., 2013; Kurudenkandy et al., 2014). Although the effects of Aβ on LTP are well established (Klyubin et al., 2012), little is known about whether and how Aβ-containing AD extracts affect pre- and post-synaptic elements. To investigate affects on presynaptic release, we measured short-term synaptic facilitation (Zucker and Regehr, 2002) in slices before and 30 min after treatment with AD extract. As synapse release probability is inversely correlated to synaptic facilitation (Zucker and Regehr, 2002), we employed high-frequency burst stimulation (5 pulses with 20 ms intra-burst stimulus interval). Application of AD extract induced a reduction in the short-term facilitation during burst stimulation (Figure 1E-F). When responses were normalized based on the ratio of each fEPSP to the first response, we found that treatment with AD extract had no effect on the 2^nd^ response, but significantly decreased the 3^rd^, 4^th^, and 5^th^ response (red dots, p= 0.02 at 3^rd^ stimulation, p= 0.004 at 4^th^ stimulation and p= 0.004 at 5^th^ stimulation, n = 6, student t-test, and also by group and time with Two way ANOVA, F(4,7)=6.39, p=0.006) (Figure 1F). In contrast, the slices treated with ID-AD yielded a pattern highly similar to that obtained with aCSF-B control (data not show). Thus, Aβ in the AD extract caused a reduction in short-term synaptic plasticity due to an initial increase in pre-synaptic glutamate release.

### Aβ-containing AD brain extract disrupts the excitation-to-inhibition balance

To estimate the effect of Aβ on the total synaptic input at the single-neuron level, we used whole-cell voltage clamp recordings to measure spontaneous excitatory postsynaptic currents (sEPSCs) on CA1 pyramidal neurons before and 30 min after addition of AD extract. The holding potential was kept constant at -70 mV and sEPSCs measured before and 30 min after addition of AD extract – this 30 min interval was chosen to match the pre-incubation time used in our LTP and short-term facilitation experiments. Application of the AD extract significantly decreased the inter-event interval (p=1.65E-6, K-S test) and increased the mean frequency of sEPSCs (from 1.8 ± 0.2 Hz to 2.7 ± 0.3 Hz, p=0.02, n = 7, students t-test) (Figure 2A and B), but did not alter the sEPSCs amplitude (mean amplitude from 11.7 ± 1.8 pA to 10.1 ± 1.6 pA, p= 0.65, n= 7, student t-test) (Fig 2A and C). In contrast, the ID-AD sample had no effect on the frequency or the amplitude of sEPSCs (mean frequency: from 2.2 ± 0.5 Hz to 2.3 ± 0.7 Hz, mean amplitude: from 9.7 ± 1.7 pA to 10.2 ± 1.4 pA, p=0.45, n = 6, student t-test) (Figure 2D–F). These results indicate that the AD brain-derived Aβ significantly augments excitatory synaptic input on CA1 pyramidal neurons.

**Figure 2.**
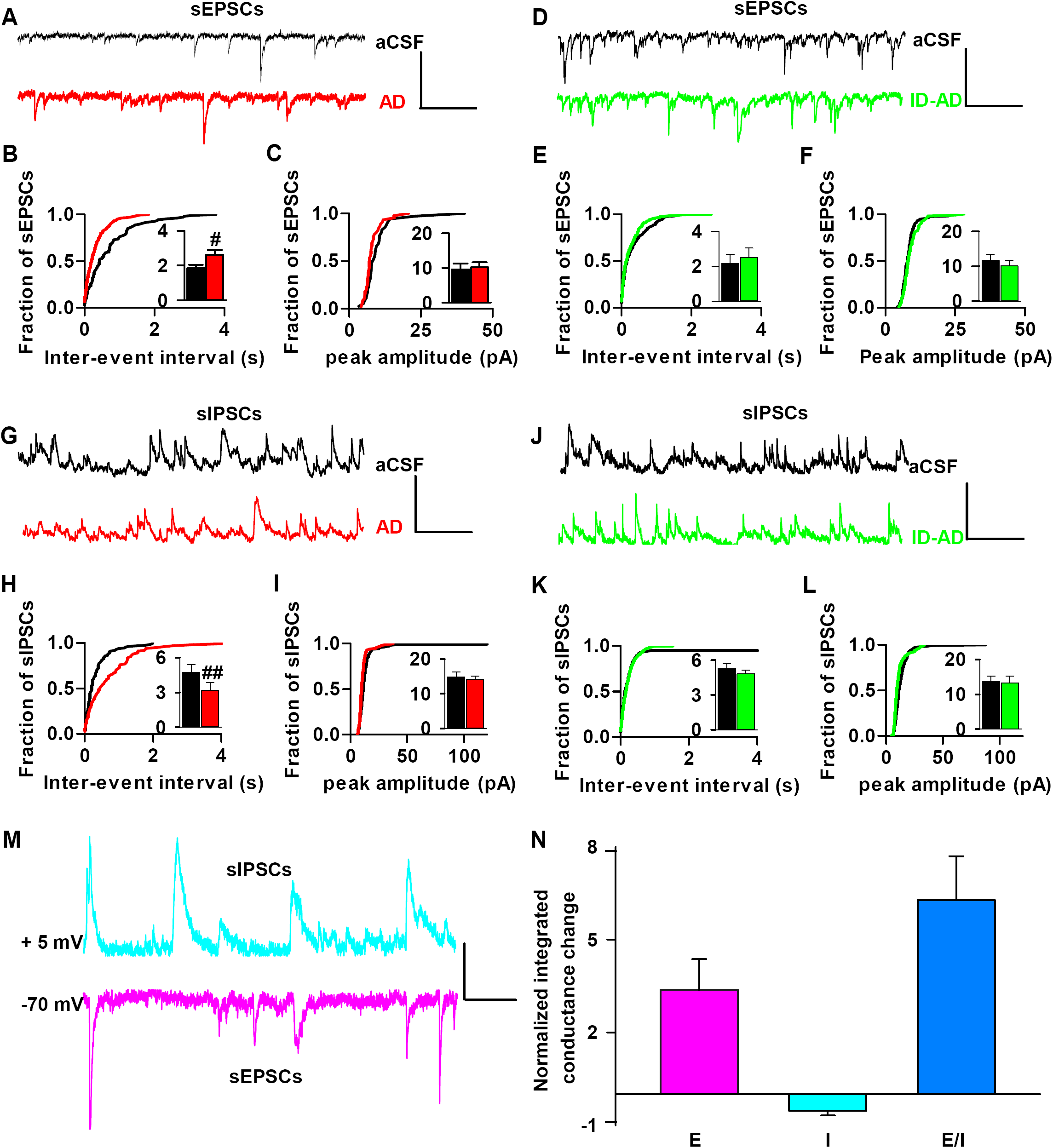
AD brain-derived Aβ affects both excitatory and inhibitory synaptic inputs, causing disruption of the excitatory/inhibitory ratio at individual CA1 neurons. (**A, D**) Example traces of spontaneous excitatory post-synaptic currents (sEPSCs) before (aCSF, black) and 30 min after addition of sample (AD, red; ID-AD, green) were recorded from individual pyramidal neurons in the hippocampal CA1 area of brain slices with the holding potential fixed at -70 mV. Scale bars: 20 pA, 700 ms. (**B**) 30 min of AD treatment decreased cumulative distributions of inter-event intervals and increased mean frequency (insert) (p=1.65E-6, K-S test; p< 0.02, student t-test; n = 7), but (**C**) did not change the cumulative distributions or the mean value (insert) of the amplitude of sEPSCs (n = 7). (**E, F**) The ID-AD sample had no effect on either frequency or amplitude of sEPSCs (n = 6). (**G, J**) Example traces of spontaneous inhibitory post-synaptic currents (sIPSCs) before (aCSF, black) and 30 min after treatment (AD, red; ID-AD, green) were recorded on the same individual pyramidal neurons upon increasing the holding potential to 5 mV. Scale bars: 20 pA, 700 ms. (**H**) 30 min of treatment with the AD sample increased inter-event intervals and decreased mean frequency (insert) of sIPSCs (red) versus aCSF (black) (p=6.19E-6, K-S test; p=0.008, student t-test; n = 7). (**I**) Treatment with the AD sample did not affect the amplitude of sIPSCs (n = 7) and the ID-AD sample had no effect on frequency (**K**) or the amplitude (**L**) of sIPSCs versus aCSF control (n = 7). **(M)** Representative traces of sIPSCs and sEPSCs from the same pyramidal neuron show charge transfer measured as the area of events above the threshold in the aCSF control. Scale bars: 10 pA, 200 ms. **(N)** Integrated conductance were measured between 30 - 35 min after addition of AD application was normalized to the value of 5 min before addition of AD. Mean excitatory integrated conductance increased and mean inhibitory integrated conductance decreased upon treatment of AD (E: excitatory input/sEPSCs; I: inhibitory input/sIPSCs). # p< 0.01.

Pyramidal neurons receive both excitatory (sEPSCs) and inhibitory (sIPSCs) inputs and GABAergic axon terminals more easily form synapses with perisomtatic regions of pyramidal cells and strongly influence the output of neurons (DeFelipe, 2002; Garcia-Marin et al., 2009). To record sIPSCs on the same neurons, we adjusted the holding potential to 5 mV, a voltage close to the calculated sEPSCs reverse potential. As shown in Figure 2G–I, the AD sample significantly increased inter-event intervals(p=6.19E-6, K-S test) and decreased the frequency of sIPSCs (from 4.7 ± 0.7 Hz to 3.1 ± 0.7 Hz, p=0.008, n = 7, student t-test), without altering sIPSCs amplitude (from 14.8 ± 1.4 pA to 14.2 ± 0.9 pA, p=0.75, n = 7, student t-test). In contrast, the ID-AD sample had no effect on sIPSCs (frequency: from 5.3 ± 0.4 Hz to 4.8 ± 0.7 Hz, amplitude: from 13.6 ± 1.6 pA to 13.2 ± 2.1 pA, p=0.21, n = 6, student t-test) (Figure 2J-L). These results revealed that brain-derived Aβ significantly reduces GABAergic input on CA1 pyramidal cells.

To assess whether the changes of excitatory input (E) and inhibitory input (I) to the same neuron affect the E/I balance of that neuron, we calculated the integrated conductance of sEPSCs and sIPSCs over a 5 min period (Figure 2M). Comparison of the charge transfer before and 30 min after AD sample application revealed E was increased ∼3 fold and I was decreased ∼50%, consequently, the E/I balance was increased ∼6 fold (n=7) (Figure 2N). These results show that AD brain-derived Aβ oppositely affects excitatory and inhibitory synaptic transmission, causing an increase in the E/I ratio. These changes, especially the reduction of GABAergic tone on individual neurons, may contribute to neuronal hyperactivity and disturb network homeostasis, and thus perturb LTP (Wang et al., 2014; Gillespie et al., 2016).

### Genetic ablation of APP occludes the effects of Aβ on LTP and pre-synaptic activity and normalizes the E/I balance

Multiple lines of evidence suggest that the APP may play a role in both GABAergic and glutamatergic neurotransmission (Bai et al., 2008; Kabogo et al., 2010; Pliassova et al., 2016; Schwenk et al., 2016) and separate studies impute a link between Aβ and APP (Lorenzo et al., 2000; Fogel et al., 2014; Kirouac et al., 2017). Thus, having found that brain-derived Aβ acts on pre-synapses and modulates both GABA and glutamate transmission, we investigated if APP was required for these effects. For this, we employed mice null for APP (Figure 3A). In agreement with prior reports, brain slices from APP KO and WT littermate mice exhibited similar levels of basal activity (P=0.19, One Way ANOVA test) and LTP (Figure 3B-F) (Dawson et al., 2000; Jedlicka et al., 2012). In both WT and APP KO slices treated with the aCSF-B control, TBS induced strong potentiation which lasted the whole recording period (158.1 ± 6.3 % in WT, n = 11, black dots; 151.2 ± 8.5 % in APP KO, n = 9, gray hexagons; F=4.4, p=0.79, comparison of the last 10 min recording using One Way ANOVA test) (Fig 3C and D). In agreement with experiments shown in Figure 1, addition of AD extract to WT slices significantly decreased LTP compared to addition of aCSF-B (121.8 ± 5.4 % in WT + AD, red dots, n = 7, F=4.5, p=0.0005, WT Ctr vs. WT + AD, One Way ANOVA test). However, application of the same extract to slices from APP KO mice had no effect on LTP, with the level of LTP in APP KOs indistinguishable from that of WT or APP KO treated with aCSF-B control (145.4 ± 4.2 % in APP KO + AD, pink upward triangles, F=4.5, p=0.41, APP KO Ctr vs. APP KO + AD; One Way ANOVA test). Similarly, AD extract had no effect on short-term facilitation (Figure 3 - figure supplement 1).

**Figure 3.**
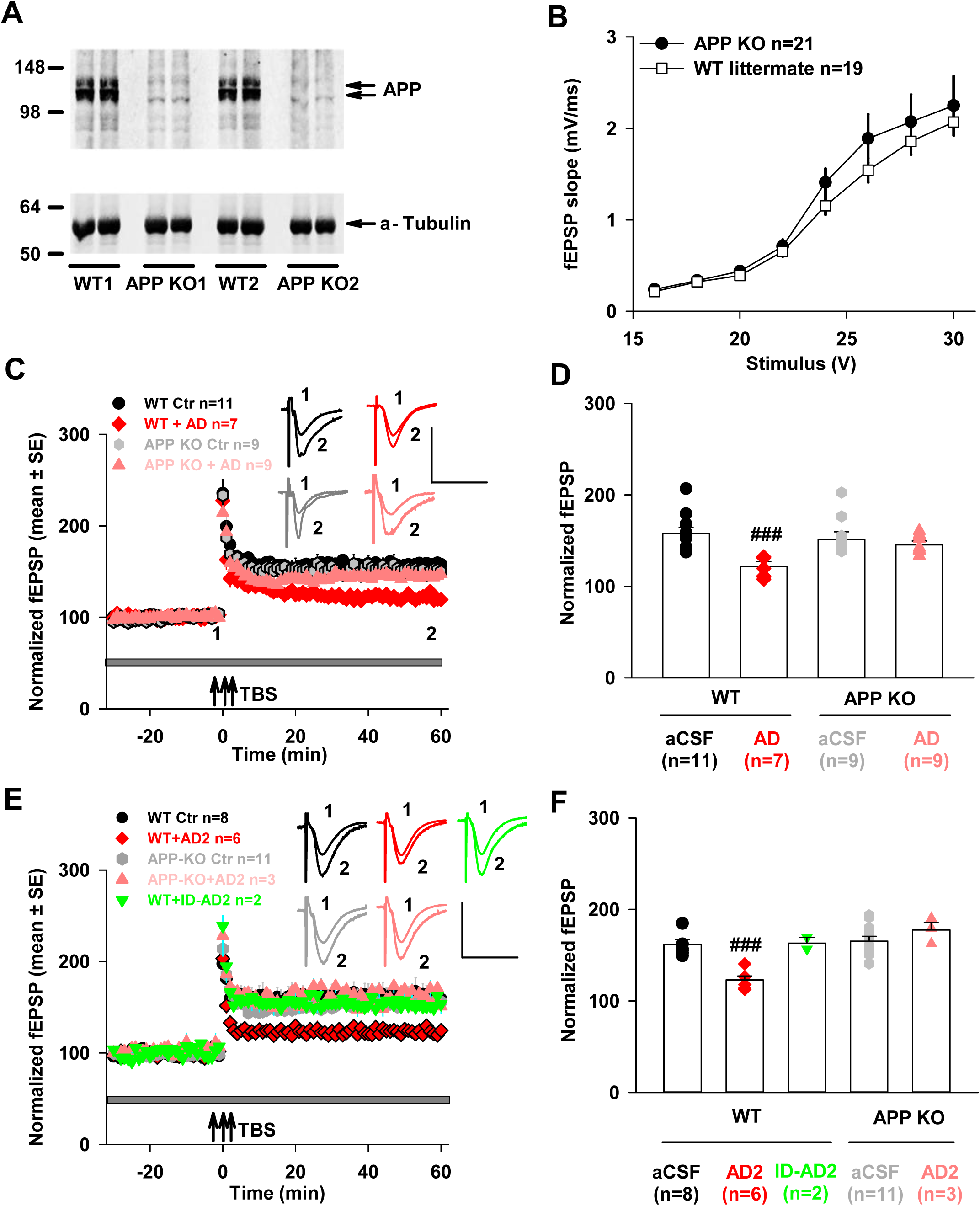
Expression of APP is required for the plasticity-disrupting activity of Aβ-containing AD brain extracts. **(A)** Aqueous extracts of mouse brain slices used for electrophysiology were analyzed for APP by Western Blotting with 22C11. Full-length APP was readily detected in extracts from wild type littermate mice (WT) but not APP knockout mice (APP KO). Slices from 2 different APP KOs (KO1 and KO2) and 2 different WTs (WT1 and WT2) are shown. **(B)** Input/output curves recorded in the hippocampal CA1 area are highly similar for both WT and APP KO mouse brain slices (p=0.19, One Way ANOVA test). Values are mean ± SEM. **(C)** LTP recorded in hippocampal CA1 was similar in brain slices from WT and APP KO mice (WT Ctr, black dots vs. APP KO Ctr, gray hexagons, p=0.79, comparison of the last 10 min recording using One Way ANOVA test). However, the extract from AD1 brain extract blocked LTP in WT but not in APP KO mice brain slices. Horizontal gray bar indicates the duration during when sample was present. 1, 2, indicate example traces from time points just prior to the theta burst stimulation (↑↑↑ TBS) (1) and 60 min after TBS (2), respectively. The aCSF control in WT mice is shown with black circles; AD treatment in WT mice is shown in red diamonds; the aCSF control in APP KO mice is shown in gray hexagons and AD treatment in APP KO mice is shown using pink upward triangles. WT slices for each treatment came from different animals; the APP KO slices came from a total of 4 APP KO mice. Scale bars: 0.5 mV, 15 ms. **(D)** Comparison of average potentiation from last 10 min of LTP recording (p F=4.5, p=0.0005, Control vs. AD in WT mice; F=4.5, p=0.41, Control vs. AD in APP KO mice; One Way ANOVA test). Symbols correspond to those in panel **C**. **(E)** Extract of a second brain (AD2) blocked hippocampal LTP in WT brain slices, but not in APP KO brain slices. Horizontal gray bar indicates the duration when sample was present. 1, 2, indicate example traces from time points just prior to the theta burst stimulation (↑↑↑ TBS) (1) and 60 min after TBS (2), respectively. Treatment of WT slices with aCSF control is shown using black circles; AD2 treatment of WT slices in red diamonds; ID-AD2 treatment in WT slices in green downward triangles. Treatment of APP KO slices with aCSF control is in gray hexagons and treatment of APP KO slices with AD2 is in pink upward triangles. WT slices for each treatment came from different animals; The APP KO slices came from a total 5 APP KO mice. Scale bars: 0.5 mV, 15 ms. **(F)** Comparison of average potentiation from last 10 min of LTP recording (F=4.8, p=0.0001, aCSF control vs. AD2 in WT mice; F=5.3, p= 0.92, ACSF control vs. ID-AD2 in WT mice; F=4.8, p=0.29, aCSF control vs. AD2 in APP KO mice; One Way ANOVA test). Symbol denoting the different treatment groups corresponds to those in panel **E**. ### p< 0.001.

To assess the generalizability of the rescue of LTP by APP ablation, we tested the effect of an extract from a second AD brain (AD2) (Figure 3 - figure supplement 2). As with the AD1 extract (Figure 1), the AD2 extract blocked LTP in slices from WT mice in an Aβ-dependent fashion (161.9 ± 5.2 % in WT Ctr, black dots, n = 8; 123 ± 4.3 % in WT + AD2, red diamonds, n = 6; F=4.8, p=0.0001, One Way ANOVA test), but had no effect on LTP elicited from APP KO mice (165.4 ± 5.2 % in APP KO Ctr, gray hexagons, n = 11; 170.5 ± 8 % in APP KO + AD2, pink upward triangles, n = 3; F=4.8, p=0.29, One Way ANOVA test) (Figure 3E and F). These results were confirmed using another APP KO line (Zheng et al., 1995) and an extract from a third AD brain (AD3) (Figure 3 - figure supplement 3). Thus, it appears that the well-documented plasticity-disrupting activity of Aβ extracted from AD brains (Klyubin et al., 2008; Shankar et al., 2008; Barry et al., 2011; Freir et al., 2011; Klyubin et al., 2012) requires expression of APP.

To investigate whether APP is necessary for the effect of Aβ on the E/I balance (Figure 2), we studied the effects of Aβ on sEPSCs and sIPSCs in brains of APP KO and WT littermate mice (Figure 4). When applied to WT slices, AD extract again increased mean sEPSC frequency (from 2.2 ± 0.1 Hz to 3.4 ± 0.2 Hz, p=0.003, n = 5, student t-test) and decreased inter-event intervals (p=6.34E-15, K-S test), without altering the amplitude of sEPSCs (mean amplitude: 17.8 ± 0.4 pA vs. 18 ± 1.5 pA, p=0.32, n = 5, student t-test) (Figure 4A-C); and on the same neuron decreased mean sIPSCs frequency (from 4.2 ± 0.8 Hz to 2.7 ± 0.4 Hz, p=0.006, n = 5, student t-test) and increased inter-event intervals (p=9.44E-20, K-S test), but not amplitude (mean amplitude from 20 ± 3 pA to 19.3 ± 1.3 pA, p=0.34, n = 5, student t-test) (Figure 4D-F). These results, which were obtained with WT mice from an entirely different colony as those used in Figure 2, nicely demonstrate the robustness of the Aβ effect (Compare Figure 2 vs. Figure 4). Most importantly, when the AD extract was applied to APP KO slices there was no change in the frequency or amplitude of sEPSCs (mean frequency: from 2.6 ± 0.1 Hz to 2.7 ± 0.4 Hz, mean amplitude: from 15 ± 1.4 pA to 14.6 ± 0.5 pA, p=0.14, K-S test; p=0.26, n = 6, student t-test) (Figure 4G-I). Similarly, sIPSCs were also unchanged (mean frequency: from 3.5 ± 0.5 Hz to 3.5 ± 0.3 Hz, mean amplitude: from 16.7 ± 1 pA to 16.4 ± 1.6 pA, p=0.58, K-S test; p=0.25, n = 6, student t-test) (Figure J-L). Thus, as with our LTP experiments (Figure 3), ablation of APP completely rescued the effects of Aβ on excitatory and inhibitory input on CA1 pyramidal neurons. Further, since APP KO occluded Aβ alterations on the E and I input at individual neurons, it also prevented Aβ-mediated changes in the integrated conductance of sEPSCs and sIPSCs (Figure 4M). When AD extract was applied to WT slices, E increased ∼3-fold and I decreased ∼44%, resulting in ∼5.8-fold increase in the E/I ratio. However APP KO significantly reversed those E/I ratio changes (p=0.001, E/I in WT vs. E/I in APP KO, One Way ANOVA test) (Figure 4M). These results indicate that APP plays an important role in regulating the acute effects of Aβ on excitatory and inhibitory pre-synaptic release, and consequent maintenance of network homeostasis.

**Figure 4.**
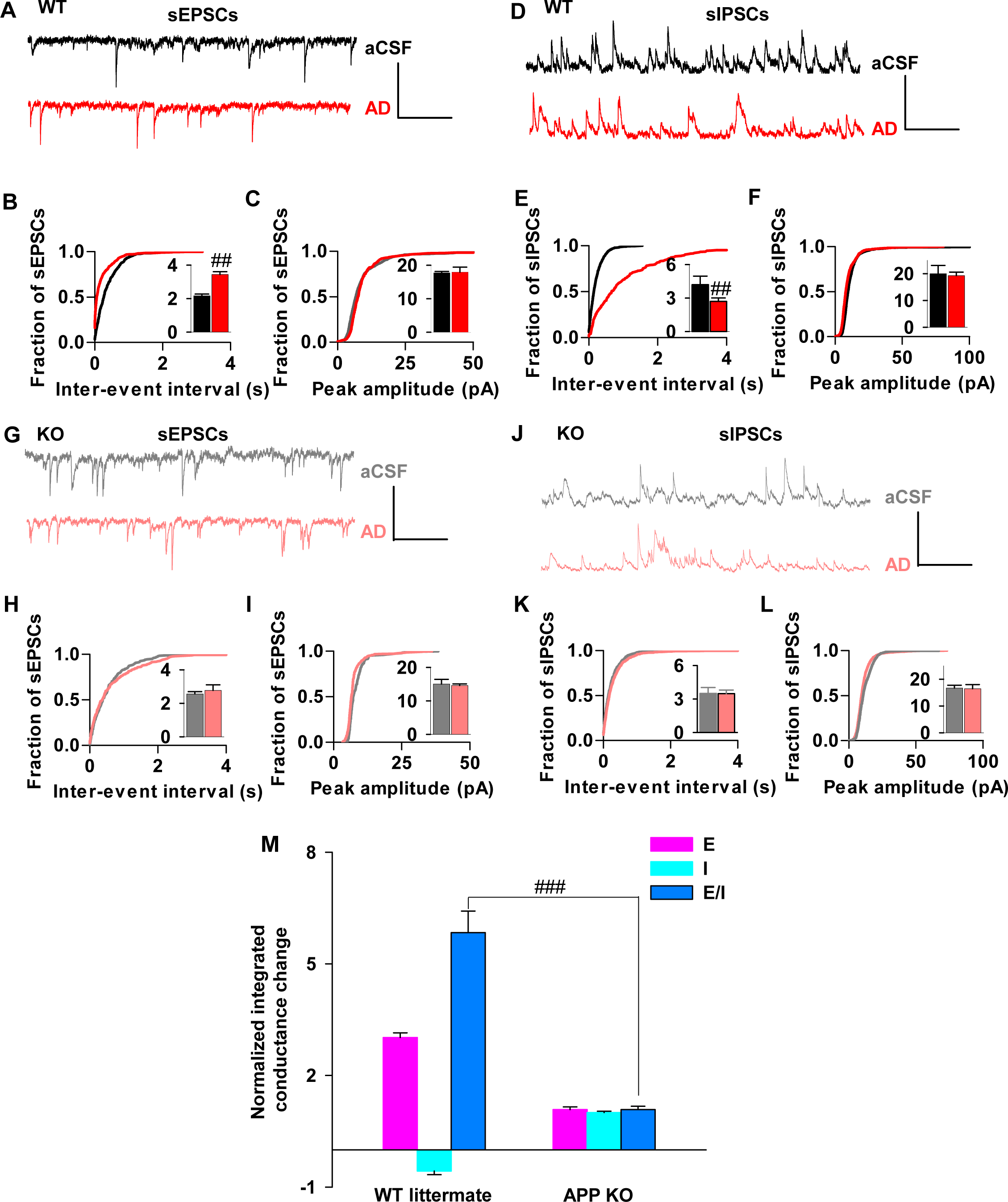
APP knock out occluded the effects of Aβ-containing AD brain extracts on both excitatory and inhibitory postsynaptic currents, and rescued the disruption of E/I balance. (**A, D**) Example traces of sEPSCs (**A**) and sIPSCs (**D**) before (aCSF, black) and 30 min after addition of AD1 extract (AD, red) on WT hippocampal brain slices. Scale bars: 20 pA, 700 ms. (**B**) Treatment with AD1 extract decreased inter-event intervals and increased mean frequency (insert) of sEPSCs (p=6.34E-15, K-S test; p=0.003, student t-test; n = 5), but (**C)** did not significantly change the cumulative distributions or the mean value (insert) of the amplitude of sEPSCs (n = 5) on WT slices. (**E**) 30 min of AD treatment increased inter-event intervals and decreased mean frequency (insert) of sIPSCs (p=9.44E-20, K-S test; p=0.006, student t-test; n = 5), but (**F)** did not affect the cumulative distributions or the mean value (insert) of the amplitude of sIPSCs (n = 5) on WT slices. (**G, J**) Example traces of spontaneous post-synaptic currents (sEPSCs, **G**; sIPSCs, **J**) before (aCSF, gray) and 30-40 min following addition of AD1 extract (AD, pink) on APP KO mice hippocampal brain slices. Scale bars: 20 pA, 700 ms. (**H**) Treatment with AD sample affected neither frequency nor amplitude (I) of sEPSCs (p=0.14, K-S test; p=0.26, student t-test; n = 6) on APP KO mice. Similarly, treatment of APP KO neurons with AD1 did not change frequency (**K**) or the amplitude (**L,** p=0.58, K-S test; p=0.25 student t-test; n = 6) of sIPSCs. (**M**) Application of Aβ-containing AD brain extract significantly changed the intergrated conductance of both excitatory (E) and inhibitory (I) input to neurons and disrupted the E/I balance in WT animals, but not in APP KO mice (p=0.001, E/I in WT vs. E/I in APP KO, One Way ANOVA test).

### Aβ binding to synapses requires APP

To further investigate the targeting of synaptic elements by Aβ and how this might be influenced by APP we used a powerful high resolution microscopic technique, array tomography (AT), to search for evidence of Aβ binding to synapses in the same brain slices used in our electrophysiology experiments. Upon completion of LTP recording, certain slices from the treatment groups used in Figs. 3C and E were immediately fixed, processed and used for AT. Brain slices were stained with 1C22, an antibody that preferentially recognizes aggregated forms of Aβ (Mably et al., 2015; Pickett et al., 2016), synapsin-1 (for pre-synapses) and PSD95 (for post synapses). Approximately 7,000 synapses (∼3,500 pre-synapses and ∼3,500 post-synapses) per slice were analyzed. AT revealed significant anti-Aβ staining at synapses of slices incubated with AD1 extract with only background staining in samples incubated with aCSF and ID controls (Figure 5A-C; Kruskal Wallis test for synapsin-1 (x^2^(4)= 10.844, p=0.028), Kruskal Wallis test for PSD95 (x^2^(4)= 11.583, p=0.021)). In slices incubated with AD1 extract 1.27 ± 0.47% of pre-synapses and 0.58 ± 0.19%of post-synapses stained with 1C22, whereas in slices that had been incubated with aCSF, only 0.0076% ± 0.013% of pre-synapses and 0.0184% ± 0.087% of post-synapses were 1C22 positive (Dunns post-hoc between AD and control for pre-synapses p=0.024 and for post-synapses p=0.010). Slices incubated with extracts immunodepeleted of Aβ exhibited similar background staining with 1C22 as the aCSF control (Figure 5A-C). Thus, the same treatment with AD1 extract that disrupts synaptic plasticity in an Aβ-dependent fashion (Figs 1 and 3) also leads to Aβ binding to synapses (Figure 5A-C). Moreover, the finding that Aβ is present at more pre-synapses than post-synapses (Mann-Whitney U between AD pre-synapses and AD post-synapses U=0, p=0.004) is consistent with our results that suggest a pre-synaptic effect of Aβ (Figure 1E and F).

**Figure 5.**
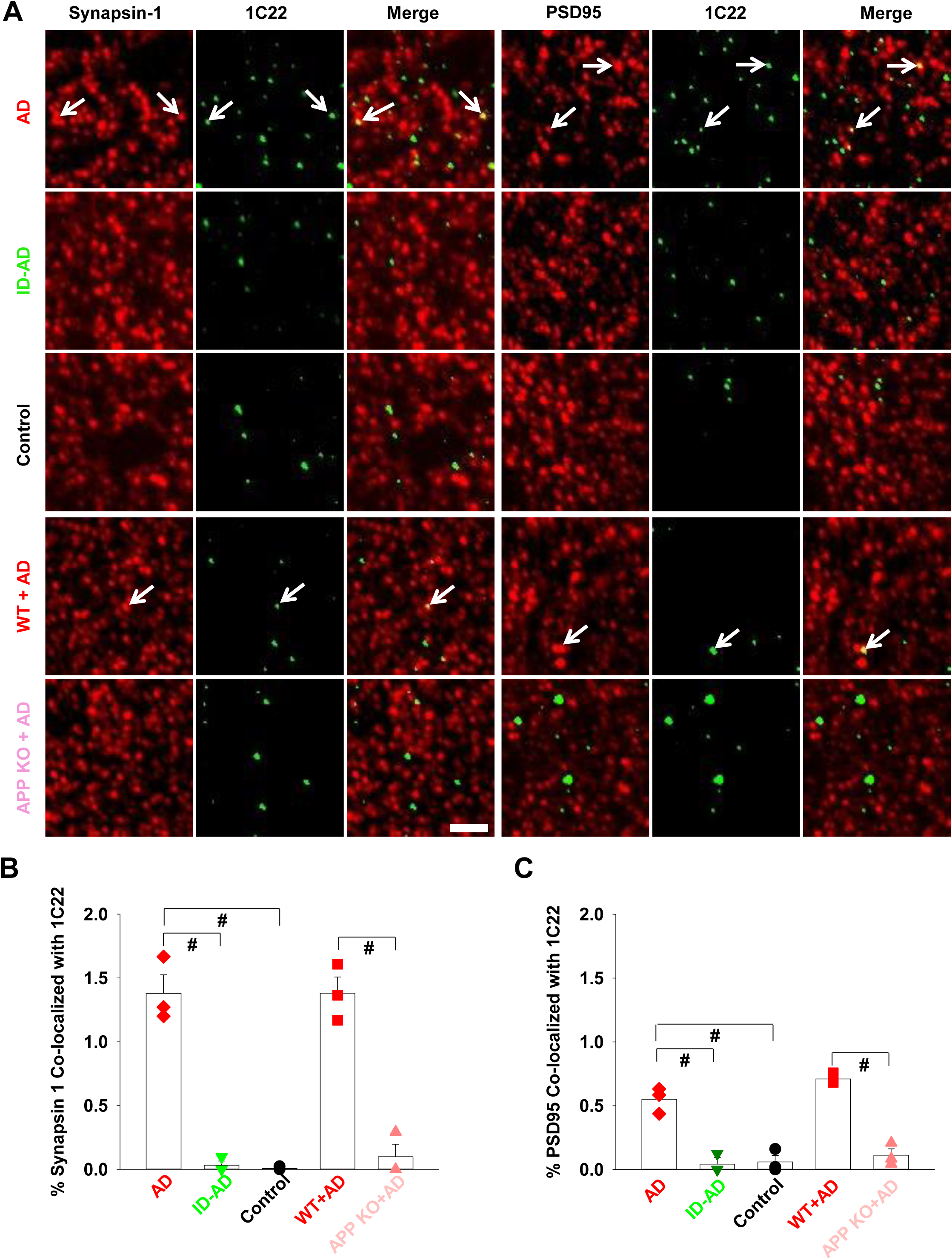
Aβ binding to synaptic terminals requires expression of APP. (**A**) Array tomography of hippocampi stained for Aβ, PSD95 (post-synapses) and synapsin-1 (pre-synapses) reveal co-localization of Aβ at synapses in slices incubated with AD brain extract. (**B** and **C**) The amount of synaptic 1C22 staining was significantly greater in slices incubated with AD extract, than in slices incubated with aCSF or ID-AD extract based on (**B**) co-localization of 1C22 and synapsin 1 staining (Kruskal Wallis test (x^2^(4)=10.844, p=0.028) Dunns post-hoc vs. control p=0.021), and (**C**) 1C22 and PSD95 co-localization (Kruskal Wallis test for PSD95 (x^2^(4)=11.583, p=0.021, Dunns post-hoc vs. control p=0.010). Importantly, when slices from APP KO mice were incubated with AD extract there was no significant co-localization of 1C22 staining with either synapsin 1 (**B**) (Dunns post-hoc vs. control p=1.000) or PSD-95 (**C**) (Dunns post-hoc vs. control p=1.000). Graphs represent the medians ± the interquartile range per treatment. Each data point is derived from the analysis of ~3,500 synapses imaged per brain slices. Within a treatment group the 3 slices used were from different mice (**B** and **C**). Arrows indicate specific examples of 1C22 staining co-localizing with pre- or post-synapses, Scale bar is 2 μm (**A**). # P< 0.05

Importantly, when brain slices from APP KO mice were incubated with AD extract, little or no synaptic 1C22 staining was detected (Figure 5A, B and C). These results are notable since expression of APP was found to be required for Aβ-mediated disruption of both long-term plasticity (Figure 3) and neurotransmitter release (Figure 4). In sum, our AT data are completely congruent with the results of our electrophysiological experiments and indicate that expression of APP is required for the binding and subsequent plasticity-disrupting effects of Aβ, and that these effects are largely mediated on the pre-synapse.

## Discussion

To better understand how Aβ disrupts synaptic plasticity we combined the use of the most disease relevant form of Aβ, Aβ extracted from human AD brain, with electrophysiological approaches and high-resolution microscopy. Consistent with prior studies, we show that extracts from the brains of individuals who died with AD block LTP (Shankar et al., 2008; Barry et al., 2011; Freir et al., 2011; Yang et al., 2017). We further show that concomitant with the block of LTP there is an increase in presynaptic release and disruption of E/I balance. In accord with these synaptic effects of Aβ, we demonstrate that exogenously applied AD brain-derived Aβ binds to synapses, with more Aβ oligomers detected on pre-synapses than on the post-synapses. Our finding that treatment with brain-derived Aβ enhances excitatory drive agrees well with studies which show that aggregated forms of synthetic Aβ increase EPSPs, action potentials, and membrane depolarizations (Hartley et al., 1999; Minkeviciene et al., 2009; Kurudenkandy et al., 2014). Our study is unique in that we employed brain-derived Aβ, and that the concentration of this material was much lower than the synthetic Aβ used in prior studies.

The apparent paradox that ectopic application of Aβ causes a net increase in excitation, yet impairs LTP may result because of glutamate spillover and activation of extra- or perisynaptic NR2B-enriched NMDARs, which play a major role in LTD induction (Li et al., 2011; Zhang et al., 2016). In such a scenario, synaptic depression may result from an initial increase in synaptic activation of NMDARs by glutamate, followed by synaptic NMDAR desensitization, NMDAR/AMPAR internalization, and activation of extrasynaptic NMDARs and mGluRs (Hu et al., 2014). However, it is not clear why ablation of APP could recover such effects.

An alternative explanation that accounts for a role for APP in the impairment of post-synaptic efficacy is that exogenous AD-derived soluble aggregates and endogenously produced monomer have differential effects. Aβ is known to be released in an activity-dependent manner (Kamenetz et al., 2003; Cirrito et al., 2005), whereas elevated Aβ levels result in depressed glutamatergic synaptic transmission and glutamate receptor endocytosis (Kamenetz et al., 2003; Hsieh et al., 2006). Thus, it is plausible that the increase in glutamate release induced by soluble Aβ aggregates may also lead to an increase in *de novo* produced Aβ monomer and this in turn may depress post-synaptic activity. Such a scenario would necessarily require expression of endogenous APP and explain why ablation of APP can obviate the block of LTP caused by brain-derived soluble Aβ aggregates. With regard to the protection of LTP upon ablation of APP, it is important to emphasize the robust nature and generalizability of this phenomenon. We observed the same protection using two different APP KO mouse lines (Zheng et al., 1995; Callahan et al., 2017) and extracts from 3 different AD brains. In all cases AD extracts blocked LTP in an Aβ-dependent manner when applied to wild type mouse brain slices, but the same AD extracts had no effect on LTP elicited from APP KO slices. Moreover, the extent of Aβ binding to synapses was similar in two different sources of wild type mice (Figure 5B and C), and the pattern observed was reminiscent of that seen in AD brain (Pickett et al., 2016).

There is evidence that APP can act as a receptor for Aβ (Melchor and Van Nostrand, 2000; Van Nostrand et al., 2002; Yankner and Lu, 2009; Fogel et al., 2014; Kirouac et al., 2017) and that APP may mediate increased - excitatory drive (Fogel et al., 2014). Specifically, Aβ was unable to promote aberrant neurotransmitter release in the absence of APP (Fogel et al., 2014). Our finding that binding of soluble Aβ aggregates to synapses requires expression of APP is consistent, but not proof, that APP may act as a receptor for Aβ. In this regard, it is worth noting that APP is known to both regulate L-type calcium channels in GABAergic neurons, interact with the pore-forming subunit Cav1.2 (Yang et al., 2009), and is a member of the GABA_B_-R receptor complex (Schwenk et al., 2016). In addition, there is evidence from proteomic studies which indicates that APP interacts with more than 30 different proteins including proteins key to synaptic vesicle turnover (Kohli et al., 2012; Del Prete et al., 2014; Lassek et al., 2014; Wilhelm et al., 2014), and proteins (such as the prion protein) which are implicated in binding Aβ (Bai et al., 2008; Lauren et al., 2009). Thus, Aβ could exert an APP-dependent effect either by directly binding to APP or binding to protein complexes of which APP is a component and stabilizing member.

So far we have considered the effects of Aβ on synapses and a single hippocampal pathway (the *Schaffer Collateral*), but Aβ is also thought to have network-wide effects (Palop and Mucke, 2010). For instance, Aβ-induced increases in excitatory network activity could lead to synaptic depression through homeostatic mechanisms. It is well established that acute treatment of primary neurons with bicuculline (a GABA_A_ antagonist) increases overall neuronal activity and firing rates (Vertkin et al., 2015). However, after a couple of days, neuronal activity returns to control levels. By analogy, it is reasonable that the disruption of E/I balance seen with our acute Aβ treatment may also cause both short-term local and long-lasting network effects. Given the fact that Aβ treatment increases excitatory drive and decreases inhibitory drive, and that GABA-ergic interneurons express high levels of APP in DG (Wang et al., 2014; Del Turco et al., 2016) it is tempting to speculate that Aβ-mediated disruption of GABA-ergic interneurons may play a special role in the cognitive impairment that occurs early in AD (Gillespie et al., 2016).

Considerable data from the study of APP tgs implicate impairment of GABAergic interneurons as central to the network disturbances evident in these models (Busche and Konnerth, 2015; Palop and Mucke, 2016). However, the unphysiological expression of high levels of APP and the concomitant release of Aβ from the expressed transgene make it difficult to differentiate between effects mediated by Aβ versus APP, or non-Aβ APP metabolites (Seabrook et al., 1999; Melnikova et al., 2013; Born et al., 2014; Fowler et al., 2014). Nonetheless, growing evidence suggests that GABAergic interneurons play a prominent role in homeostatic regulation of hippocampal networks and there is compelling proteomic and physiological data that link APP and GABA_B1a_-R (Wang et al., 2014; Gillespie et al., 2016; Schwenk et al., 2016). Consequently further investigations on how Aβ effects GABA_B_-R expression, GABA_B_-R-APP interactions and whether GABA_B_-R KOs are resistant to Aβ are merited and may lead to a pharmacological means to attenuate Aβ synaptotoxicity. Similarly, modulation of APP expression may also offer therapeutic potential. However, while our results demonstrate that ablation of APP in brain slices from young (2-3 month) mice protects against the acute synaptotoxicity of Aβ, widespread knock-out of APP is not recommended. APP appears to be involved in many physiological processes (Yang et al., 2009; Muller and Zheng, 2012; Del Prete et al., 2014; Lassek et al., 2014; Wang et al., 2014) and aged APP null mice exhibit hypersensitivity to kainate-induced seizures (Steinbach et al., 1998), altered exploratory behavior, deficits in spatial memory, and impairment of LTP (Dawson et al., 1999; Phinney et al., 1999; Seabrook et al., 1999; Ring et al., 2007). No such deficits have been reported in APP hemizygous mice, thus it maybe possible to down regulate APP expression so as to maintain normal function, yet attenuate Aβ synaptotoxicity.

## Materials and Methods

### Reagents

All chemicals and reagents were purchased from Sigma-Aldrich unless otherwise noted. Synthetic Aβ1–42 was synthesized and purified using reversed-phase HPLC by Dr. James I. Elliott at the ERI Amyloid laboratory Oxford, CT, USA. Peptide mass and purity (>99%) were confirmed by reversed-phase HPLC and electrospray/ion trap mass spectrometry.

### Antibodies

The antibodies used and their source are described in Table 1.

**Table 1.** Primary and secondary antibodies.

| Antibody | Type | Antigen/epitope | Dilution for IP | Conc. For WB | Conc. For ELISA | Dilution for AT | Source/Reference |
| --- | --- | --- | --- | --- | --- | --- | --- |
| 3D6 | Monoclonal | Aβ1–5 | – | - | 1 μg/ml | – | Elan/(Johnson-Wood et al., 1997) |
| 266 | Monoclonal | Aβ16-23 | – | - | 3 μg/ml | – | Elan/(Seubert et al., 1992) |
| 2G3 | Monoclonal | Aβ terminating at Val40 | – | 1 μg/ml | – | – | Elan/(Johnson-Wood et al., 1997) |
| 21F12 | Monoclonal | Aβ terminating at Ile42 | – | 1 μg/ml | 1 μg/ml | – | Elan/(Johnson-Wood et al., 1997) |
| 1C22 | Monoclonal | Aβ aggregates | – | – | 3 μg/ml | 1:50 | Walsh lab/(Mably et al., 2015) |
| AW7 | Polyclonal | Pan anti-Aβ | 1:80 | – | – | – | Walsh lab/(Mc Donald et al., 2015) |
| 22C11 | Monoclonal | APP66-81 | – | 1 μg/ml | – | 1:50 | Millipore |
| AB1543 P | Polyclonal | Rabbit anti-synapsin-1 | – | – | – | 1:100 | Millipore/(Kay et al., 2013) |
| 3450P | Polyclonal | Rabbit anti-PSD95 | – | – | – | 1:5 | Cell Signaling/(Kay et al., 2013) |
| A21202 | Polyclonal | Donkey anti-mouse 488 | – | – | – | 1:50 | Invitrogen |
| A21207 | Polyclonal | Donkey anti-rabbit 594 | – | – | – | 1:50 | Invitrogen |

### Preparation of human brain extracts

All human specimens were obtained and used in accordance with the Partner’s Institutional Review Board (Protocol: Walsh BWH 2011). Tissue was from the brains of individuals (referred to as AD1, AD2 and AD3) who died with AD. AD1 was an 87 years old man who 9 months prior to death had scored 23 on the MMSE and at postmortem had pathological changes consistent with mild AD. AD2 was a 65 years old female who 3 years prior to death scored 24 on the MMSE and at postmortem was diagnosed as having AD. AD3 was a 68 years old female with end-stage AD and fulminant amyloid and neurofibrillary tangles pathology. Aqueous extracts of brain were prepared by homogenizing cortical tissue in a buffer which we refer to as artificial cerebrospinal fluid base buffer (aCSF-B) (124 mM NaCl, 2.8 mM KCl, 1.25 mM NaH_2_PO_4_, 26 mM NaHCO_3_, pH 7.4). aCSF-B is the core buffer used in subsequent electrophysiology experiments. Whole frozen temporal cortex was left at 4°C until the tissue was sufficiently soft to cut. Meninges and large blood vessels were removed and gray matter dissected from white matter. The total amount of gray matter obtained was between 12-14 g. Two gram lots of tissue were diced using a razor blade and then homogenized in 10 ml of ice-cold aCSF-B (containing 5 mM Ethylenediaminetetraacetic acid, 1 mM Ethyleneglycoltetraacetic acid, 5 μg/ml Leupeptin, 5 μg/ml Aprotinin, 2 μg/ml Pepstatin, 120 μg/ml Pefabloc and 5 mM NaF) with 25 strokes of a Dounce homogenizer (Fisher, Ottawa, Canada). Homogenates from 6, 2 g lots were pooled and centrifuged at 198,000 g and 4°C for 110 min in a SW 41 Ti rotor (Beckman Coulter, Fullerton, CA). The upper 90% of supernatant was dialyzed (using Slide-A-Lyzer™ G2 Dialysis Cassettes, 2K MWCO, Fisher Scientific) against fresh aCSF-B to remove bioactive small molecules and drugs. Dialysis was performed at 4°C against a 100-fold excess of buffer with buffer changed 3 times over a 36 h period. Thereafter extracts were divided into 2 parts: 1 portion was immunodepleted (ID) of Aβ by 3 rounds of 12 hour incubations with the anti-Aβ antibody, AW7, plus Protein A sepharose (PAS) beads at 4°C (Freir et al., 2011). The second portion was treated in an identical manner, but this time incubated with pre-immune serum plus PAS beads. Samples were cleared of beads and 0.5 ml aliquots stored at -80°C until used for biochemical or electrophysiological experiments. Samples were thawed once and used.

### Immunoprecipitation/Western blotting (IP/WB) of Aβ in brain extracts

Extracts were first pre-cleared with PAS beads to minimize non-specific interactions in the subsequent IP. One ml aliquots of extracts were incubated with 15 μl PAS beads for 1 hour at 4°C with gentle shaking. Aβ-antibody-PAS beads were removed by centrifugation (4000 g for 5 minutes) and the supernatant divided into 0.5 ml aliquots. Each aliquot was incubated with 10 μl of AW7 and 15 1/4l PAS beads overnight at 4°C with gentle shaking. PAS complexes were collected by centrifugation and washed as previously described (Shankar et al., 2011). The immunoprecipitated (IP’d) Aβ was eluted by boiling in 18 1/4l of 1 × sample buffer (50 mM Tris, 2% w/v SDS, 12% v/v glycerol with 0.01% phenol red) and electrophoresed on hand poured, 15 well 16% polyacrylamide tris-tricine gels. Synthetic Aβ1-42 was run as a loading control and protein transferred onto 0.2 μM nitrocellulose at 400 mA and 4°C for 2 h. Blots were microwaved in PBS and Aβ detected using the anti-Aβ40 and anti-Aβ42 antibodies, 2G3 and 21F12, and bands visualized using a Li-COR Odyssey infrared imaging system (Li-COR, Lincoln, NE).

### MSD Aβ immunoassays

Samples were analyzed for Aβ content using 2 distinct assay formats: the Aβx-42 assay that preferentially detects Aβ42 monomers and the oAssay that preferentially detects Aβ oligomers and aggregates (Mably et al., 2015; Mc Donald et al., 2015; Yang et al., 2015). Immunoassays were performed using the Meso Scale Discovery (MSD) platform and reagents from Meso Scale (Rockville, MD). The Aβx-42 assay uses mAb m266 (3 μg/ml) for capture and biotinylated 21F12 (1 μg/ml) for detection, and the oAssay uses mAb 1C22 (3 μg/ml) for capture and biotinylated 3D6 (1 μg/ml) for detection. Samples, standards and blanks were loaded in triplicate and analyzed as described previously (Mc Donald et al., 2015).

Since GuHCl effectively disaggregates high molecular weight Aβ species (Mc Donald et al., 2015), samples were analyzed both with and without incubation in 5 M GuHCl. Analysis of samples in the absence of GuHCl allows the measurement of native Aβ42 monomer using the Aβx-42 assay, and native Aβ aggregates using the oAssay. Analysis of samples treated with GuHCl allows detection of disassembled aggregates with Aβx-42 assay. To dissociate aggregates 20 μl of extract was incubated overnight with 50 μl of 7 M GuHCl at 4°C. Thereafter samples were diluted 1:10 with assay diluent, so that the final GuHCl concentration was 0.5 M. Aβ standards were prepared in tris-buffered saline (TBS), pH 7.4 containing 0.5 M GuHCl, 0.05% Tween 20 and 1% Blocker A so that both standards and samples contained the same final concentration of GuHCl.

### Mice

All animal procedures were performed in accordance with the National Institutes of Health Policy on the Use of Animals in Research and were approved by the Harvard Medical School Standing Committee on Animals. Wild type (WT) C57BL/6 mice were purchased from Jackson Labs (Bar Harbor, ME). APP KO mice on a C57BL/6 background and littermate WT controls were obtained from the Young-Pearse lab (Callahan et al., 2017). A second line of APP KO mice were purchased from the Jackson Laboratory (APP^tm1Dbo^/J,The Jackson Laboratory, Bar Harbor, ME) (Zheng et al., 1995). Animals were housed in a room with a 12 h light/dark circadian cycle with *ad libitum* access to food and water. Mice were genotyped by PCR prior to use, and after use certain brain slices were used for Western blotting (Figure 3A).

### Brain slices preparation

Two to three months old male and female animals were anaesthetized with isoflurane and decapitated. Brains were rapidly removed and immediately immersed in ice-cold (0-4°C) artificial cerebrospinal fluid (aCSF). The aCSF contained (in mM): 124 NaCl, 3 KCl, 2.4 CaCl_2_, 2 MgSO_4_·7H_2_O, 1.25 NaH_2_PO_4_, 26 NaHCO_3_ and 10 D-glucose, and was equilibrated with 95% O_2_ and 5% CO_2_, pH 7.4, 310 mOsm. Coronal brain slices (350μm) including hippocampus (Wang et al., 2008) were prepared using a Leica VT1000 S vibratome (Leica Biosystems Inc, Buffalo Grove, IL) and transferred to an interface chamber and incubated at 34 ± 5°C for 20 min and then kept at room temperature for 1 h before recording.

### Long-term potentiation (LTP) recording

Brian slices were transferred to a submerged recording chamber and perfused (10 ml/min) with oxygenated (95% O_2_ and 5% CO_2_) aCSF 10 min before electrophysiological recording. Brian slices were visualized using an infrared and differential interference contrast camera (IR-DIC camera, Hitachi, Japan) mounted on an upright Olympus microscope (Olympus, Tokyo, Japan). Recording electrodes were pulled from borosilicate glass capillaries (Sutter Instruments, Novato, CA) using a micropipette puller (Model P-97; Sutter Instruments, Novato, CA) with resistance ∼2 MΩ when filled with ACSF. To induce field excitatory post-synaptic potential (fEPSP) in the hippocampal CA1, a tungsten wire stimulating electrode (FHC, Inc., Bowdoin, ME) was placed on the Schaffer collaterals of the CA3 and a recording electrode was placed at least 300 μM away on the striatum radiatum of the CA1. Test stimuli were delivered once every 20 s (0.05 Hz) and the stimulus intensity was adjusted to produce a baseline fEPSP of 30–40% of the maximal response. A stable baseline was recorded for at least 10 min prior to addition of sample. Thirty minutes following application of sample LTP was induced by theta burst stimulation (TBS, involved 3 trains, each of 4 pulses delivered at 100 Hz, 10 times, with an interburst interval of 200 ms with a 20 sec interval between each train). Field potentials were recorded using a Multiclamp amplifier (Multiclamp 700B; Molecular Devices, Sunnyvale, CA) coupled to a Digidata 1440A digitizer. Signal was sampled at 10 kHz and filtered at 2 kHz and data were analyzed using Clampex 10 software (Molecular Devices, Sunnyvale, CA).

### Whole-cell patch clamp recording

Brain slices were prepared from male and female WT and APP KO mice (1-2 months old) as described above for LTP experiments but using a different cutting solution contained sucrose (in mM: 72 sucrose, 83 NaCl, 2.5 KCl, 1 NaH_2_PO_4_, 3.3 MgSO4·7H2O, 26.2 NaHCO_3_, 22 dextrose, and 0.5 CaCl_2_) saturated with 95% O_2_ and 5% CO_2_, pH 7.4, 310 mOsm (Wang et al., 2015). Slices were incubated in oxygenated slicing solution for 20 min, and held at room temperature for a further 40 min prior to recording. Slices were transferred to a submerged recording chamber and perfused (10 ml/min) with oxygenated (95% O_2_ and 5% CO_2_) aCSF for 30 min at room temperature. Whole-cell recordings were made from the somata of CA1 pyramidal neurons visualized using an IR-DIC camera mounted on an upright Olympus microscope (Olympus, Tokyo, Japan). Patch pipettes (4–7MΩ) were filled with an internal solution containing (in mM): 120 CsGluconate, 5 MgCl_2_, 0.6 EGTA, 30 HEPES, 4 MgATP, 0.4 Na_2_GTP, 10 phosphocreatine-Tris, 5 QX-314; 290 mOsm; pH was adjusted at 7.2 with CsOH. Signal was acquired using a Multiclamp amplifier (Multiclamp 700B; Molecular Devices, Sunnyvale, CA) with Clampex 10 software (Molecular Devices, Sunnyvale, CA) and sampled at 10 kHz and filtered at 2 kHz. Data were stored on a PC after digitization by an A/D converter (Digidata 1440A, Molecular Devices, Sunnyvale, CA) for offline analysis. Membrane potential was corrected for the liquid junction potential of 13.7 mV. Neurons with negative resting membrane potential less than -60 mV were not analyzed. Input resistance and patching access resistance were continuously monitored during the experiment and cells which exhibited more than 15–20% changes in these parameters were excluded from analysis.

In order to preserve a relatively intact neuronal circuit no receptor antagonists were used. Spontaneous excitatory post-synaptic currents (sEPSCs) were collected at a membrane holding potential of -70 mV, which is close to the calculated reverse potential of GABA. In order to measure the excitatory and inhibitory input on the same neuron, the spontaneous inhibitory post-synaptic currents (sIPSCs) were also measured on the same neuron but this time the holding potential was increased to 5-10 mV, a potential close to the reverse potential of excitatory input, without visual negative deflection. Recorded neuronal activities were detected as described previously (Lillis et al., 2015) by custom software (DClamp: available at www.ieeg.org/?q=node/34). Integrated excitatory conductance (sEPSCs, G_E_) and intergreated inhibitory conductance (sIPSCs,G_I_) were calculated as previously reported 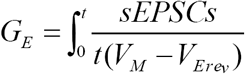 and 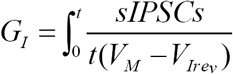 (Slomowitz et al., 2015).

### Preparation of mouse brain homogenates and detection of APP

Certain brain slices from wild-type and APP knock-out mice were frozen immediately after completion of electrophysiological recording (Figs 3 and 4) and stored at -80°C until analyzed. Tissue (∼0.1 mg) was homogenized in 5 volumes (w/v) of ice-cold 20 mM TBS-TX, pH 7.4 containing protease inhibitors and centrifuged at 100,000 g and 4°C for 78 minutes in a TLA-55 rotor (Beckman Coulter, Fullerton, CA). The upper 90% of the supernatant was removed, aliquoted and stored at -80°C pending analysis. Ten μg of total protein was boiled in 1 × sample buffer (62.5 mM Tris, 1% w/v SDS, 10% v/v glycerol, 0.01% phenol red and 2% β-mercaptoethanol) for 5 min and then electrophoresed on hand poured, 15 well 10% polyacrylamide tris-glycine gels. Gels were rinsed in transfer buffer (10% methanol, 192 mM Glycine and 25 mM Tris) and proteins electroblotted onto 0.2 μM nitrocellulose membranes at 400 mA and 4°C for 2.5 h. Membranes were developed using the anti-APP antibody, 22C11, and bands visualized using a LI-COR Odyssey infrared imaging system (LI-COR, Lincoln, NE).

### Array tomography (AT) imaging of mouse brain slices

Upon completion of electrophysiology recordings certain brain slices from wild-type and APP knock mice (Figs 3 and 4) were processed for array tomography (Koffie et al., 2009; Pickett et al., 2016). Slices were fixed in PBS containing 4% paraformaldehyde and 2.5% sucrose at 4°C overnight. Samples were then washed three times (10 min each) in cold wash buffer (PBS containing 3.5% sucrose and 50 mM glycine), and the hippocampus dissected out under a Leica Wild M3Z Stereozoom Microscope (Heerbrugg, Swizerland). Thereafter hippocampi were dehydrated using an ethanol series of: 50%, 70%, 95% and 100%. Tissue was then placed into a solution of 1:1 ethanol: LR White resin (Electron Microscopy Sciences) for 5 min and then washed 3 times with LR white. Tissue was incubated overnight at 4°C in LR white and then embedded in a gelatin capsule and polymerized overnight at 53°C. Three embedded blocks per condition were cut into ribbons of 70 nm sections on an ultracut microtome (Leica) using a Jumbo Histo Diamond Knife (Diatome). Ribbons were collected on gelatin-coated glass coverslips, stained with antibodies and imaged along the ribbon. Two ribbons per slice were collected and one was stained for PSD95 and 1C22 and the other for synapsin-1 and 1C22. Primary antibodies were 1C22 (1:50), rabbit anti-PSD95 (3450P, Cell Signaling, at 1:5), and rabbit anti-synapsin-1 (AB1543P, Millipore, at 1:100). Secondary antibodies donkey anti-mouse 488 (A21202) and donkey anti-rabbit 594 (A21207) were from Invitrogen and used at 1:50.

Two image stacks per ribbon were collected from the stratum radiatum using a Zeiss axio Imager Z2 epifluorescent microscope with a 63X 1.4NA Plan Apochromat objective. Images were acquired with a CoolSnap digital camera and AxioImager software with array tomography macros (Carl Zeiss, Ltd, Cambridge UK). Images from each set of serial sections were complied to create a 3D stack and aligned using ImageJ multistackreg macros (Kay et al., 2013). Regions of interest (10 μm x 10 μm) were selected, cropped and thresholded in Image J (Schindelin et al., 2012; Ollion et al., 2013). Custom MatLab macros were used to remove single slice punctuate, count synaptic punctuate and assess co-localization with 1C22 (a minimum of 50% overlap between 1C22 and synaptic punctuate was required to be designated as co-localization). All custom analysis macros will be freely available on http://datashare.is.ed.ac.uk after publication.

### Data analysis and Statistics test

IP/WB and MSD Aβ immunoassay is representative of at least 2 experiments. For electrophysiologcial experiments, the AD, ID-AD and aCSF samples were coded and tested in an interleaved manner to avoid variances in animals or slice quality influencing on results. There was no outliners were excluded from the analysis. Electrophysioloical data were analyzed offline by pclamp 10.2 (Molecular Devices, Sunnyvale, CA) and tested with One-way or Two-way analysis of variance (ANOVA) with Bonferroni post-hoc tests or student t-tests (# P<0.05, ## P<0.01, and ### P<0.001). A Kolmogorov–Smirnov (K–S) test was used to compute differences in distributions of sEPSCs and sIPSCs. Array tomography was analyzed using SPSS Version 22. A single percent co-localization for each parameter was calculated for each slice from approximately 41 regions of interest and ≈7500 synapses (∼3,500 pre-synapses and ∼3,500 post-synapses) were analyzed per slice and tested with a Kruskal-Wallis with Dunns post-hoc test and Mann-Whitney U test. Electrophysiology data are shown as means ± SEM. Array tomography data is shown as medians ± the interquartile range, each point representing all synapses measured within 1 slice. Analyses of the same sample using different slices are considered technical replicates and analysis of extracts from different AD brains are considered biological replicates.

---

**Figure 3 – figure supplement 1:**
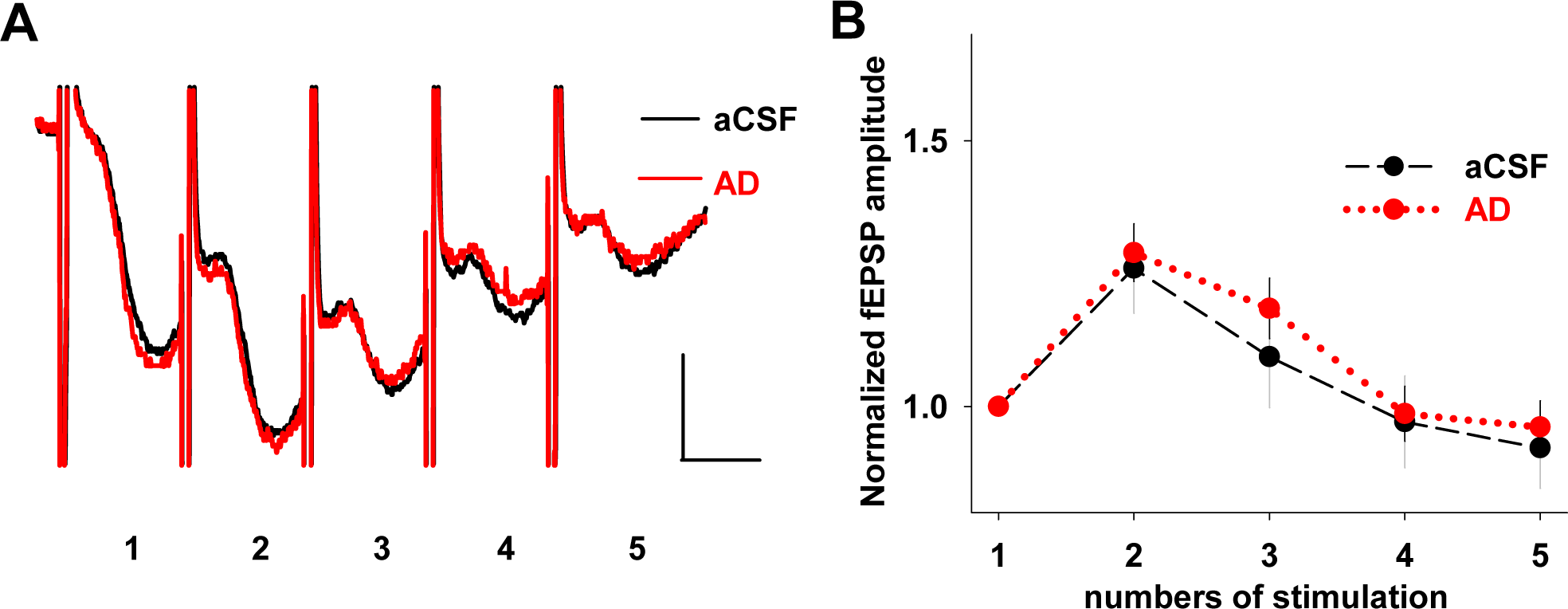
APP ablation occludes the effect of AD brain extract on short term facilitation. (**A**) Representative traces of averaged field recordings were collected after 5 stimulation bursts (inter-stimulation interval 20 ms, inter-burst interval 30 s) before (black, aCSF) and 30 min after perfusion with the AD sample (red) on brain slice from APP KO mouse. Scale bars: 0.5 mV, 10 ms. (**B**) fEPSPs amplitude after 2 to 5 stimulations were normalized to the value obtained after the first stimulation. Values are mean ± SEM. There is no significant difference between aCSF control and the presence of AD brain extract application (n=5, F=5.32, P=0.7, One Way ANOVA test).

**Figure 3 – figure supplement 2:**
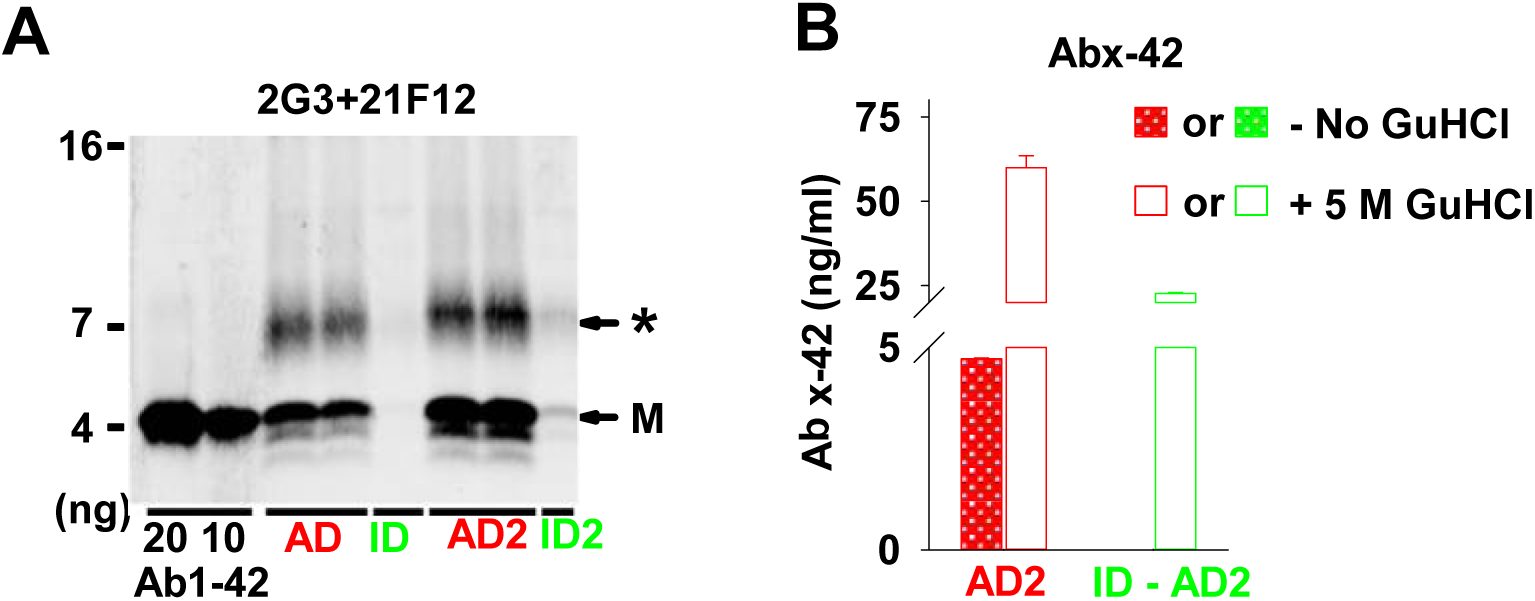
Characterization of Aβ in the water-soluble extract from AD2 brain. (**A**) Aqueous extract of AD1 and AD2 were treated with either pre-immune serum or AW7 antiserum. Portions of the mock immunodepleted sample (AD1 and AD2, red) and the AW7 immunodepleted sample (ID-AD1 and ID-AD2, green) were then analyzed by IP/WB, using AW7 for IP and a combination of 2G3 and 21F12 for WB. **M** denotes Aβ monomer and * indicates a broad smear ~7–8 kDa. No specific bands were detected above 16 kDa marker and the blot was cropped accordingly. (**B)** AD2 (red) and ID-AD2 (green) samples were incubated +/- 5 M GuHCl and analyzed using an immunoassay that preferentially recognize Aβx-42 monomer (266-21F12b). AW7 ID reduced the monomer signal from 4.73 ± 0.02 ng/ml to undetectable levels. Upon GuHCl treatment, 60 ± 3.52 ng/ml of Aβ x-42 was detected in AD2 and AW7 ID decreased this to 22.72 ± 0.7 ng/ml.

**Figure 3 – figure supplement 3:**
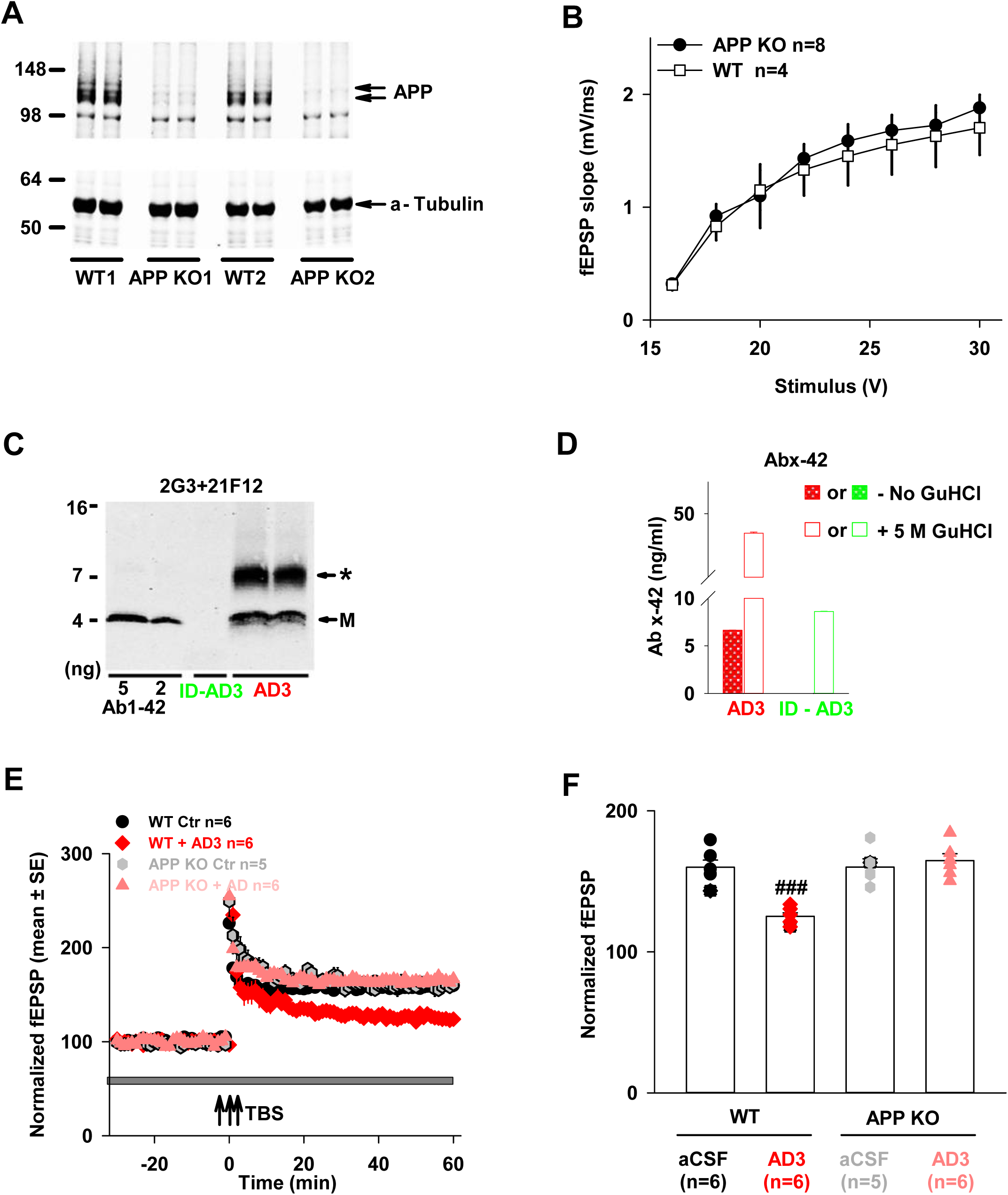
APP ablation occludes the plasticity-disrupting activity of AD3 brain extract. **(A)** Wild type (WT) and APP knock-out (KO) mouse brain slices used for electrophysiology were analyzed for APP by Western Blotting with 22C11. Full-length APP was readily detected in extracts from WT but not APP KO. Slices from 2 different KOs (KO1 and KO2) and WTs (WT1 and WT2) mice are shown. **(B)** Input/output curves recorded in the hippocampal CA1 area are highly similar for both WT and APP KO mouse brain slices (F=4.6, p= 0.73, One Way ANOVA test). Values are mean ± SEs. **(C)** Aqueous extract of AD3 was treated with either pre-immune serum or with AW7 antiserum. Portions of the mock immunodepleted sample (AD3, red) and the AW7 immunodepleted sample (ID-AD3, green) were then analyzed by IP/WB, using AW7 for IP and a combination of 2G3 and 21F12 for WB. **M** denotes Aβ monomer and * indicates a broad smear ~7–8 kDa. No specific bands were detected above 16 kDa marker and the blot was cropped accordingly. (**D)** AD3 (red) and ID-AD3 (green) samples were incubated +/- 5 M GuHCl and analyzed using an immunoassay that preferentially recognize Aβ42 monomer (266-21F12b). AW7 ID reduced monomer from 6.65 ± 0.01 ng/ml to undetectable level without GuHCl treatment. Upon treatment with GuHCl, the amount of Aβ42 increased to 46.94 ± 0.2 ng/ml in AD3 and this was reduced to 8.62 ± 0.1 ng/ml by immunodepletion. **(E)** LTP recorded in hippocampal CA1 was similar in brain slices from WT and APP KO mice. Notably, the extract from AD3 blocked LTP in WT but not in APP KO mice brain slices. Horizontal gray bar indicates the duration during when sample was present. The aCSF control in WT mice is shown with black circles; AD treatment in WT mice is shown in red diamonds; the aCSF control in APP KO mice is shown in gray hexagons and AD treatment in APP KO mice is shown using pink upward triangles. WT slices for each treatment came from different animals; the APP KO slices came from a total of 4 APP KO mice. Theta burst stimulation (↑↑↑ TBS). Scale bars: 0.5 mV, 15 ms. **(F)** Comparison of average potentiation from last 10 min of LTP recording (F=4.96, p= 0.0001, Control vs. AD in WT mice; F=5.12, p= 0.56, Control vs. AD in APP KO mice; One Way ANOVA test). Symbols correspond to those in panel **E**.

## References

Abramov E, Dolev I, Fogel H, Ciccotosto GD, Ruff E, Slutsky I (2009) Amyloid-beta as a positive endogenous regulator of release probability at hippocampal synapses. Nature neuroscience 12:1567–1576.

Ashe KH, Zahs KR (2010) Probing the biology of Alzheimer’s disease in mice. Neuron 66:631–645.

Bai Y, Markham K, Chen F, Weerasekera R, Watts J, Horne P, Wakutani Y, Bagshaw R, Mathews PM, Fraser PE, Westaway D, St George-Hyslop P, Schmitt-Ulms G (2008) The in vivo brain interactome of the amyloid precursor protein. Molecular & cellular proteomics: MCP 7:15–34.

Barry AE, Klyubin I, Mc Donald JM, Mably AJ, Farrell MA, Scott M, Walsh DM, Rowan MJ (2011) Alzheimer’s disease brain-derived amyloid-beta-mediated inhibition of LTP in vivo is prevented by immunotargeting cellular prion protein. The Journal of neuroscience: the official journal of the Society for Neuroscience 31:7259–7263.

Borlikova GG, Trejo M, Mably AJ, Mc Donald JM, Sala Frigerio C, Regan CM, Murphy KJ, Masliah E, Walsh DM (2013) Alzheimer brain-derived amyloid beta-protein impairs synaptic remodeling and memory consolidation. Neurobiology of aging 34:1315–1327.

Born HA, Kim JY, Savjani RR, Das P, Dabaghian YA, Guo Q, Yoo JW, Schuler DR, Cirrito JR, Zheng H, Golde TE, Noebels JL, Jankowsky JL (2014) Genetic suppression of transgenic APP rescues Hypersynchronous network activity in a mouse model of Alzeimer’s disease. The Journal of neuroscience: the official journal of the Society for Neuroscience 34:3826–3840.

Busche MA, Konnerth A (2015) Neuronal hyperactivity–A key defect in Alzheimer’s disease? BioEssays: news and reviews in molecular, cellular and developmental biology 37:624–632.

Callahan DG, Taylor WM, Tilearcio M, Cavanaugh T, Selkoe DJ, Young-Pearse TL (2017) Embryonic mosaic deletion of APP results in displaced Reelin-expressing cells in the cerebral cortex. Developmental biology 424:138–146.

Cirrito JR, Yamada KA, Finn MB, Sloviter RS, Bales KR, May PC, Schoepp DD, Paul SM, Mennerick S, Holtzman DM (2005) Synaptic activity regulates interstitial fluid amyloid-beta levels in vivo. Neuron 48:913–922.

Cleary JP, Walsh DM, Hofmeister JJ, Shankar GM, Kuskowski MA, Selkoe DJ, Ashe KH (2005) Natural oligomers of the amyloid-beta protein specifically disrupt cognitive function. Nature neuroscience 8:79–84.

Conboy L, Murphy KJ, Regan CM (2005) Amyloid precursor protein expression in the rat hippocampal dentate gyrus modulates during memory consolidation. Journal of neurochemistry 95:1677-1688.

Dawson GR, Seabrook GR, Zheng H, Smith DW, Graham S, O'Dowd G, Bowery BJ, Boyce S, Trumbauer ME, Chen HY, Van der Ploeg LH, Sirinathsinghji DJ (1999) Age-related cognitive deficits, impaired long-term potentiation and reduction in synaptic marker density in mice lacking the beta-amyloid precursor protein. Neuroscience 90:1–13.

De Strooper B (2010) Proteases and proteolysis in Alzheimer disease: a multifactorial view on the disease process. Physiological reviews 90:465–494.

DeFelipe J (2002) Cortical interneurons: from Cajal to 2001. Progress in brain research 136:215–238.

Del Prete D, Lombino F, Liu X, D'Adamio L (2014) APP is cleaved by Bace1 in pre-synaptic vesicles and establishes a pre-synaptic interactome, via its intracellular domain, with molecular complexes that regulate pre-synaptic vesicles functions. PloS one 9:e108576.

Del Turco D, Paul MH, Schlaudraff J, Hick M, Endres K, Muller UC, Deller T (2016) Region-Specific Differences in Amyloid Precursor Protein Expression in the Mouse Hippocampus. Frontiers in molecular neuroscience 9:134.

Doyle E, Bruce MT, Breen KC, Smith DC, Anderton B, Regan CM (1990) Intraventricular infusions of antibodies to amyloid-beta-protein precursor impair the acquisition of a passive avoidance response in the rat. Neuroscience letters 115:97–102.

Esch FS, Keim PS, Beattie EC, Blacher RW, Culwell AR, Oltersdorf T, McClure D, Ward PJ (1990) Cleavage of amyloid beta peptide during constitutive processing of its precursor. Science 248:1122–1124.

Fanutza T, Del Prete D, Ford MJ, Castillo PE, D'Adamio L (2015) APP and APLP2 interact with the synaptic release machinery and facilitate transmitter release at hippocampal synapses. eLife 4:e09743.

Fogel H, Frere S, Segev O, Bharill S, Shapira I, Gazit N, O'Malley T, Slomowitz E, Berdichevsky Y, Walsh DM, Isacoff EY, Hirsch JA, Slutsky I (2014) APP homodimers transduce an amyloid-beta-mediated increase in release probability at excitatory synapses. Cell reports 7:1560–1576.

Fowler SW, Chiang AC, Savjani RR, Larson ME, Sherman MA, Schuler DR, Cirrito JR, Lesne SE, Jankowsky JL (2014) Genetic modulation of soluble Abeta rescues cognitive and synaptic impairment in a mouse model of Alzheimer’s disease. The Journal of neuroscience: the official journal of the Society for Neuroscience 34:7871–7885.

Freir DB, Nicoll AJ, Klyubin I, Panico S, Mc Donald JM, Risse E, Asante EA, Farrow MA, Sessions RB, Saibil HR, Clarke AR, Rowan MJ, Walsh DM, Collinge J (2011) Interaction between prion protein and toxic amyloid beta assemblies can be therapeutically targeted at multiple sites. Nature communications 2:336.

Garcia-Marin V, Blazquez-Llorca L, Rodriguez JR, Boluda S, Muntane G, Ferrer I, Defelipe J (2009) Diminished perisomatic GABAergic terminals on cortical neurons adjacent to amyloid plaques. Frontiers in neuroanatomy 3:28.

Gillespie AK, Jones EA, Lin YH, Karlsson MP, Kay K, Yoon SY, Tong LM, Nova P, Carr JS, Frank LM, Huang Y (2016) Apolipoprotein E4 Causes Age-Dependent Disruption of Slow Gamma Oscillations during Hippocampal Sharp-Wave Ripples. Neuron 90:740–751.

Golde TE, Estus S, Younkin LH, Selkoe DJ, Younkin SG (1992) Processing of the amyloid protein precursor to potentially amyloidogenic derivatives. Science 255:728–730.

Guerreiro R, Hardy J (2014) Genetics of Alzheimer’s disease. Neurotherapeutics: the journal of the American Society for Experimental NeuroTherapeutics 11:732–737.

Haass C, Schlossmacher MG, Hung AY, Vigo-Pelfrey C, Mellon A, Ostaszewski BL, Lieberburg I, Koo EH, Schenk D, Teplow DB, et al. (1992) Amyloid beta-peptide is produced by cultured cells during normal metabolism. Nature 359:322–325.

Hartley DM, Walsh DM, Ye CP, Diehl T, Vasquez S, Vassilev PM, Teplow DB, Selkoe DJ (1999) Protofibrillar intermediates of amyloid beta-protein induce acute electrophysiological changes and progressive neurotoxicity in cortical neurons. The Journal of neuroscience: the official journal of the Society for Neuroscience 19:8876–8884.

Hsieh H, Boehm J, Sato C, Iwatsubo T, Tomita T, Sisodia S, Malinow R (2006) AMPAR removal underlies Abeta-induced synaptic depression and dendritic spine loss. Neuron 52:831–843.

Hu NW, Nicoll AJ, Zhang D, Mably AJ, O'Malley T, Purro SA, Terry C, Collinge J, Walsh DM, Rowan MJ (2014) mGlu5 receptors and cellular prion protein mediate amyloid-beta-facilitated synaptic long-term depression in vivo. Nature communications 5:3374.

Huang JK, Ma PL, Ji SY, Zhao XL, Tan JX, Sun XJ, Huang FD (2013) Age-dependent alterations in the presynaptic active zone in a Drosophila model of Alzheimer’s disease. Neurobiology of disease 51:161–167.

Huber G, Martin JR, Loffler J, Moreau JL (1993) Involvement of amyloid precursor protein in memory formation in the rat: an indirect antibody approach. Brain research 603:348–352.

Johnson KA, Fox NC, Sperling RA, Klunk WE (2012) Brain imaging in Alzheimer disease. Cold Spring Harbor perspectives in medicine 2:a006213.

Kabogo D, Rauw G, Amritraj A, Baker G, Kar S (2010) ss-amyloid-related peptides potentiate K+-evoked glutamate release from adult rat hippocampal slices. Neurobiology of aging 31:1164–1172.

Kamenetz F, Tomita T, Hsieh H, Seabrook G, Borchelt D, Iwatsubo T, Sisodia S, Malinow R (2003) APP processing and synaptic function. Neuron 37:925–937.

Kay KR, Smith C, Wright AK, Serrano-Pozo A, Pooler AM, Koffie R, Bastin ME, Bak TH, Abrahams S, Kopeikina KJ, McGuone D, Frosch MP, Gillingwater TH, Hyman BT, Spires-Jones TL (2013) Studying synapses in human brain with array tomography and electron microscopy. Nature protocols 8:1366–1380.

Kim J, Chakrabarty P, Hanna A, March A, Dickson DW, Borchelt DR, Golde T, Janus C (2013) Normal cognition in transgenic BRI2-Abeta mice. Molecular neurodegeneration 8:15.

Kirouac L, Rajic AJ, Cribbs DH, Padmanabhan J (2017) Activation of Ras-ERK Signaling and GSK-3 by Amyloid Precursor Protein and Amyloid Beta Facilitates Neurodegeneration in Alzheimer’s Disease. eNeuro 4.

Klyubin I, Cullen WK, Hu NW, Rowan MJ (2012) Alzheimer’s disease Abeta assemblies mediating rapid disruption of synaptic plasticity and memory. Molecular brain 5:25.

Klyubin I, Betts V, Welzel AT, Blennow K, Zetterberg H, Wallin A, Lemere CA, Cullen WK, Peng Y, Wisniewski T, Selkoe DJ, Anwyl R, Walsh DM, Rowan MJ (2008) Amyloid beta protein dimer-containing human CSF disrupts synaptic plasticity: prevention by systemic passive immunization. The Journal of neuroscience: the official journal of the Society for Neuroscience 28:4231–4237.

Koffie RM, Meyer-Luehmann M, Hashimoto T, Adams KW, Mielke ML, Garcia-Alloza M, Micheva KD, Smith SJ, Kim ML, Lee VM, Hyman BT, Spires-Jones TL (2009) Oligomeric amyloid beta associates with postsynaptic densities and correlates with excitatory synapse loss near senile plaques. Proceedings of the National Academy of Sciences of the United States of America 106:4012– 4017.

Kohli BM, Pflieger D, Mueller LN, Carbonetti G, Aebersold R, Nitsch RM, Konietzko U (2012) Interactome of the amyloid precursor protein APP in brain reveals a protein network involved in synaptic vesicle turnover and a close association with Synaptotagmin-1. Journal of proteome research 11:4075–4090.

Kurudenkandy FR, Zilberter M, Biverstal H, Presto J, Honcharenko D, Stromberg R, Johansson J, Winblad B, Fisahn A (2014) Amyloid-beta-induced action potential desynchronization and degradation of hippocampal gamma oscillations is prevented by interference with peptide conformation change and aggregation. The Journal of neuroscience: the official journal of the Society for Neuroscience 34:11416–11425.

Lambert MP, Barlow AK, Chromy BA, Edwards C, Freed R, Liosatos M, Morgan TE, Rozovsky I, Trommer B, Viola KL, Wals P, Zhang C, Finch CE, Krafft GA, Klein WL (1998) Diffusible, nonfibrillar ligands derived from Abeta1-42 are potent central nervous system neurotoxins. Proceedings of the National Academy of Sciences of the United States of America 95:6448–6453.

Lassek M, Weingarten J, Einsfelder U, Brendel P, Muller U, Volknandt W (2013) Amyloid precursor proteins are constituents of the presynaptic active zone. Journal of neurochemistry 127:48–56.

Lassek M, Weingarten J, Acker-Palmer A, Bajjalieh SM, Muller U, Volknandt W (2014) Amyloid precursor protein knockout diminishes synaptic vesicle proteins at the presynaptic active zone in mouse brain. Current Alzheimer research 11:971–980.

Lassek M, Weingarten J, Wegner M, Mueller BF, Rohmer M, Baeumlisberger D, Arrey TN, Hick M, Ackermann J, Acker-Palmer A, Koch I, Muller U, Karas M, Volknandt W (2016) APP Is a Context-Sensitive Regulator of the Hippocampal Presynaptic Active Zone. PLoS computational biology 12:e1004832.

Lauren J, Gimbel DA, Nygaard HB, Gilbert JW, Strittmatter SM (2009) Cellular prion protein mediates impairment of synaptic plasticity by amyloid-beta oligomers. Nature 457:1128–1132.

Li S, Hong S, Shepardson NE, Walsh DM, Shankar GM, Selkoe D (2009) Soluble oligomers of amyloid Beta protein facilitate hippocampal long-term depression by disrupting neuronal glutamate uptake. Neuron 62:788–801.

Li S, Jin M, Koeglsperger T, Shepardson NE, Shankar GM, Selkoe DJ (2011) Soluble Abeta oligomers inhibit long-term potentiation through a mechanism involving excessive activation of extrasynaptic NR2B-containing NMDA receptors. The Journal of neuroscience: the official journal of the Society for Neuroscience 31:6627–6638.

Lillis KP, Wang Z, Mail M, Zhao GQ, Berdichevsky Y, Bacskai B, Staley KJ (2015) Evolution of Network Synchronization during Early Epileptogenesis Parallels Synaptic Circuit Alterations. The Journal of neuroscience: the official journal of the Society for Neuroscience 35:9920–9934.

Lorenzo A, Yuan M, Zhang Z, Paganetti PA, Sturchler-Pierrat C, Staufenbiel M, Mautino J, Vigo FS, Sommer B, Yankner BA (2000) Amyloid beta interacts with the amyloid precursor protein: a potential toxic mechanism in Alzheimer’s disease. Nature neuroscience 3:460–464.

Mably AJ, Kanmert D, Mc Donald JM, Liu W, Caldarone BJ, Lemere CA, O'Nuallain B, Kosik KS, Walsh DM (2015) Tau immunization: a cautionary tale? Neurobiology of aging 36:1316–1332.

Mc Donald JM, O'Malley TT, Liu W, Mably AJ, Brinkmalm G, Portelius E, Wittbold WM, 3rd, Frosch MP, Walsh DM (2015) The aqueous phase of Alzheimer’s disease brain contains assemblies built from approximately 4 and approximately 7 kDa Abeta species. Alzheimer’s & dementia: the journal of the Alzheimer’s Association 11:1286–1305.

Melchor JP, Van Nostrand WE (2000) Fibrillar amyloid beta-protein mediates the pathologic accumulation of its secreted precursor in human cerebrovascular smooth muscle cells. The Journal of biological chemistry 275:9782–9791.

Melnikova T, Fromholt S, Kim H, Lee D, Xu G, Price A, Moore BD, Golde TE, Felsenstein KM, Savonenko A, Borchelt DR (2013) Reversible pathologic and cognitive phenotypes in an inducible model of Alzheimer-amyloidosis. The Journal of neuroscience: the official journal of the Society for Neuroscience 33:3765–3779.

Mileusnic R, Lancashire CL, Johnston AN, Rose SP (2000) APP is required during an early phase of memory formation. The European journal of neuroscience 12:4487–4495.

Minkeviciene R, Rheims S, Dobszay MB, Zilberter M, Hartikainen J, Fulop L, Penke B, Zilberter Y, Harkany T, Pitkanen A, Tanila H (2009) Amyloid beta-induced neuronal hyperexcitability triggers progressive epilepsy. The Journal of neuroscience: the official journal of the Society for Neuroscience 29:3453–3462.

Mockett BG, Richter M, Abraham WC, Muller UC (2017) Therapeutic Potential of Secreted Amyloid Precursor Protein APPsalpha. Frontiers in molecular neuroscience 10:30.

Muller UC, Zheng H (2012) Physiological functions of APP family proteins. Cold Spring Harbor perspectives in medicine 2:a006288.

Neve RL, McPhie DL (2007) Dysfunction of amyloid precursor protein signaling in neurons leads to DNA synthesis and apoptosis. Biochimica et biophysica acta 1772:430–437.

Nilsson P, Saito T, Saido TC (2014) New mouse model of Alzheimer’s. ACS chemical neuroscience 5:499– 502.

Nimmrich V, Grimm C, Draguhn A, Barghorn S, Lehmann A, Schoemaker H, Hillen H, Gross G, Ebert U, Bruehl C (2008) Amyloid beta oligomers (A beta(1-42) globulomer) suppress spontaneous synaptic activity by inhibition of P/Q-type calcium currents. The Journal of neuroscience: the official journal of the Society for Neuroscience 28:788–797.

Ollion J, Cochennec J, Loll F, Escude C, Boudier T (2013) TANGO: a generic tool for high-throughput 3D image analysis for studying nuclear organization. Bioinformatics 29:1840–1841.

Palop JJ, Mucke L (2009) Epilepsy and cognitive impairments in Alzheimer disease. Archives of neurology 66:435–440.

Palop JJ, Mucke L (2010) Amyloid-beta-induced neuronal dysfunction in Alzheimer’s disease: from synapses toward neural networks. Nature neuroscience 13:812–818.

Palop JJ, Mucke L (2016) Network abnormalities and interneuron dysfunction in Alzheimer disease. Nature reviews Neuroscience 17:777–792.

Parodi J, Sepulveda FJ, Roa J, Opazo C, Inestrosa NC, Aguayo LG (2010) Beta-amyloid causes depletion of synaptic vesicles leading to neurotransmission failure. The Journal of biological chemistry 285:2506–2514.

Phinney AL, Deller T, Stalder M, Calhoun ME, Frotscher M, Sommer B, Staufenbiel M, Jucker M (1999) Cerebral amyloid induces aberrant axonal sprouting and ectopic terminal formation in amyloid precursor protein transgenic mice. The Journal of neuroscience: the official journal of the Society for Neuroscience 19:8552–8559.

Pickett EK, Koffie RM, Wegmann S, Henstridge CM, Herrmann AG, Colom-Cadena M, Lleo A, Kay KR, Vaught M, Soberman R, Walsh DM, Hyman BT, Spires-Jones TL (2016) Non-Fibrillar Oligomeric Amyloid-beta within Synapses. Journal of Alzheimer’s disease: JAD 53:787–800.

Pliassova A, Lopes JP, Lemos C, Oliveira CR, Cunha RA, Agostinho P (2016) The Association of Amyloid-beta Protein Precursor With alpha-and beta-Secretases in Mouse Cerebral Cortex Synapses Is Altered in Early Alzheimer’s Disease. Molecular neurobiology 53:5710–5721.

Portelius E, Olsson M, Brinkmalm G, Ruetschi U, Mattsson N, Andreasson U, Gobom J, Brinkmalm A, Holtta M, Blennow K, Zetterberg H (2013) Mass spectrometric characterization of amyloid-beta species in the 7PA2 cell model of Alzheimer’s disease. Journal of Alzheimer’s disease: JAD 33:85– 93.

Ring S, Weyer SW, Kilian SB, Waldron E, Pietrzik CU, Filippov MA, Herms J, Buchholz C, Eckman CB, Korte M, Wolfer DP, Muller UC (2007) The secreted beta-amyloid precursor protein ectodomain APPs alpha is sufficient to rescue the anatomical, behavioral, and electrophysiological abnormalities of APP-deficient mice. The Journal of neuroscience: the official journal of the Society for Neuroscience 27:7817–7826.

Ripoli C, Piacentini R, Riccardi E, Leone L, Li Puma DD, Bitan G, Grassi C (2013) Effects of different amyloid beta-protein analogues on synaptic function. Neurobiology of aging 34:1032–1044.

Russell CL, Semerdjieva S, Empson RM, Austen BM, Beesley PW, Alifragis P (2012) Amyloid-beta acts as a regulator of neurotransmitter release disrupting the interaction between synaptophysin and VAMP2. PloS one 7:e43201.

Scheff SW, Price DA, Schmitt FA, Mufson EJ (2006) Hippocampal synaptic loss in early Alzheimer’s disease and mild cognitive impairment. Neurobiology of aging 27:1372–1384.

Scheff SW, Price DA, Schmitt FA, DeKosky ST, Mufson EJ (2007) Synaptic alterations in CA1 in mild Alzheimer disease and mild cognitive impairment. Neurology 68:1501–1508.

Schindelin J, Arganda-Carreras I, Frise E, Kaynig V, Longair M, Pietzsch T, Preibisch S, Rueden C, Saalfeld S, Schmid B, Tinevez JY, White DJ, Hartenstein V, Eliceiri K, Tomancak P, Cardona A (2012) Fiji: an open-source platform for biological-image analysis. Nature methods 9:676–682.

Schwenk J, Perez-Garci E, Schneider A, Kollewe A, Gauthier-Kemper A, Fritzius T, Raveh A, Dinamarca MC, Hanuschkin A, Bildl W, Klingauf J, Gassmann M, Schulte U, Bettler B, Fakler B (2016) Modular composition and dynamics of native GABAB receptors identified by high-resolution proteomics. Nature neuroscience 19:233–242.

Seabrook GR, Smith DW, Bowery BJ, Easter A, Reynolds T, Fitzjohn SM, Morton RA, Zheng H, Dawson GR, Sirinathsinghji DJ, Davies CH, Collingridge GL, Hill RG (1999) Mechanisms contributing to the deficits in hippocampal synaptic plasticity in mice lacking amyloid precursor protein. Neuropharmacology 38:349–359.

Shaked GM, Kummer MP, Lu DC, Galvan V, Bredesen DE, Koo EH (2006) Abeta induces cell death by direct interaction with its cognate extracellular domain on APP (APP 597-624). FASEB journal: official publication of the Federation of American Societies for Experimental Biology 20:1254– 1256.

Shankar GM, Walsh DM (2009) Alzheimer’s disease: synaptic dysfunction and Abeta. Molecular neurodegeneration 4:48.

Shankar GM, Welzel AT, McDonald JM, Selkoe DJ, Walsh DM (2011) Isolation of low-n amyloid beta-protein oligomers from cultured cells, CSF, and brain. Methods Mol Biol 670:33–44.

Shankar GM, Li S, Mehta TH, Garcia-Munoz A, Shepardson NE, Smith I, Brett FM, Farrell MA, Rowan MJ, Lemere CA, Regan CM, Walsh DM, Sabatini BL, Selkoe DJ (2008) Amyloid-beta protein dimers isolated directly from Alzheimer’s brains impair synaptic plasticity and memory. Nature medicine 14:837–842.

Sisodia SS (1992) Beta-amyloid precursor protein cleavage by a membrane-bound protease. Proceedings of the National Academy of Sciences of the United States of America 89:6075–6079.

Soba P, Eggert S, Wagner K, Zentgraf H, Siehl K, Kreger S, Lower A, Langer A, Merdes G, Paro R, Masters CL, Muller U, Kins S, Beyreuther K (2005) Homo- and heterodimerization of APP family members promotes intercellular adhesion. The EMBO journal 24:3624–3634.

Sokolow S, Luu SH, Nandy K, Miller CA, Vinters HV, Poon WW, Gylys KH (2012) Preferential accumulation of amyloid-beta in presynaptic glutamatergic terminals (VGluT1 and VGluT2) in Alzheimer’s disease cortex. Neurobiology of disease 45:381–387.

Sola Vigo F, Kedikian G, Heredia L, Heredia F, Anel AD, Rosa AL, Lorenzo A (2009) Amyloid-beta precursor protein mediates neuronal toxicity of amyloid beta through Go protein activation. Neurobiology of aging 30:1379–1392.

Steinbach JP, Muller U, Leist M, Li ZW, Nicotera P, Aguzzi A (1998) Hypersensitivity to seizures in beta-amyloid precursor protein deficient mice. Cell death and differentiation 5:858–866.

Tamayev R, Matsuda S, Arancio O, D'Adamio L (2012) beta-but not gamma-secretase proteolysis of APP causes synaptic and memory deficits in a mouse model of dementia. EMBO molecular medicine 4:171–179.

Tanzi RE (2012) The genetics of Alzheimer disease. Cold Spring Harbor perspectives in medicine 2.

Van Nostrand WE, Melchor JP, Keane DM, Saporito-Irwin SM, Romanov G, Davis J, Xu F (2002) Localization of a fibrillar amyloid beta-protein binding domain on its precursor. The Journal of biological chemistry 277:36392–36398.

Vertkin I, Styr B, Slomowitz E, Ofir N, Shapira I, Berner D, Fedorova T, Laviv T, Barak-Broner N, Greitzer-Antes D, Gassmann M, Bettler B, Lotan I, Slutsky I (2015) GABAB receptor deficiency causes failure of neuronal homeostasis in hippocampal networks. Proceedings of the National Academy of Sciences of the United States of America 112:E3291–3299.

Walsh DM, Teplow DB (2012) Alzheimer’s disease and the amyloid beta-protein. Progress in molecular biology and translational science 107:101–124.

Walsh DM, Klyubin I, Fadeeva JV, Cullen WK, Anwyl R, Wolfe MS, Rowan MJ, Selkoe DJ (2002) Naturally secreted oligomers of amyloid beta protein potently inhibit hippocampal long-term potentiation in vivo. Nature 416:535–539.

Wang B, Wang Z, Sun L, Yang L, Li H, Cole AL, Rodriguez-Rivera J, Lu HC, Zheng H (2014) The amyloid precursor protein controls adult hippocampal neurogenesis through GABAergic interneurons. The Journal of neuroscience: the official journal of the Society for Neuroscience 34:13314– 13325.

Wang ZM, Qi YJ, Wu PY, Zhu Y, Dong YL, Cheng ZX, Zhu YH, Dong Y, Ma L, Zheng P (2008) Neuroactive steroid pregnenolone sulphate inhibits long-term potentiation via activation of alpha2-adrenoreceptors at excitatory synapses in rat medial prefrontal cortex. The international journal of neuropsychopharmacology 11:611–624.

Weidemann A, Konig G, Bunke D, Fischer P, Salbaum JM, Masters CL, Beyreuther K (1989) Identification, biogenesis, and localization of precursors of Alzheimer’s disease A4 amyloid protein. Cell 57:115– 126.

Welzel AT, Maggio JE, Shankar GM, Walker DE, Ostaszewski BL, Li S, Klyubin I, Rowan MJ, Seubert P, Walsh DM, Selkoe DJ (2014) Secreted amyloid beta-proteins in a cell culture model include N-terminally extended peptides that impair synaptic plasticity. Biochemistry 53:3908–3921.

White AR, Zheng H, Galatis D, Maher F, Hesse L, Multhaup G, Beyreuther K, Masters CL, Cappai R (1998) Survival of cultured neurons from amyloid precursor protein knock-out mice against Alzheimer’s amyloid-beta toxicity and oxidative stress. The Journal of neuroscience: the official journal of the Society for Neuroscience 18:6207–6217.

Wilhelm BG, Mandad S, Truckenbrodt S, Krohnert K, Schafer C, Rammner B, Koo SJ, Classen GA, Krauss M, Haucke V, Urlaub H, Rizzoli SO (2014) Composition of isolated synaptic boutons reveals the amounts of vesicle trafficking proteins. Science 344:1023–1028.

Willem M et al. (2015) eta-Secretase processing of APP inhibits neuronal activity in the hippocampus. Nature 526:443–447.

Yang L, Wang Z, Wang B, Justice NJ, Zheng H (2009) Amyloid precursor protein regulates Cav1.2 L-type calcium channel levels and function to influence GABAergic short-term plasticity. The Journal of neuroscience: the official journal of the Society for Neuroscience 29:15660–15668.

Yang T, Li S, Xu H, Walsh DM, Selkoe DJ (2017) Large Soluble Oligomers of Amyloid beta-Protein from Alzheimer Brain Are Far Less Neuroactive Than the Smaller Oligomers to Which They Dissociate. The Journal of neuroscience: the official journal of the Society for Neuroscience 37:152–163.

Yang T, O'Malley TT, Kanmert D, Jerecic J, Zieske LR, Zetterberg H, Hyman BT, Walsh DM, Selkoe DJ (2015) A highly sensitive novel immunoassay specifically detects low levels of soluble Abeta oligomers in human cerebrospinal fluid. Alzheimer’s research & therapy 7:14.

Yankner BA, Lu T (2009) Amyloid beta-protein toxicity and the pathogenesis of Alzheimer disease. The Journal of biological chemistry 284:4755–4759.

Zhang D, Mably AJ, Walsh DM, Rowan MJ (2016) Peripheral Interventions Enhancing Brain Glutamate Homeostasis Relieve Amyloid beta- and TNFalpha-Mediated Synaptic Plasticity Disruption in the Rat Hippocampus. Cerebral cortex.

Zheng H, Jiang M, Trumbauer ME, Sirinathsinghji DJ, Hopkins R, Smith DW, Heavens RP, Dawson GR, Boyce S, Conner MW, Stevens KA, Slunt HH, Sisoda SS, Chen HY, Van der Ploeg LH (1995) beta-Amyloid precursor protein-deficient mice show reactive gliosis and decreased locomotor activity. Cell 81:525–531.

Zucker RS, Regehr WG (2002) Short-term synaptic plasticity. Annual review of physiology 64:355–405.

